# Structural Insights into Sphingosine-1-phosphate Receptor Activation

**DOI:** 10.1101/2022.01.15.475352

**Authors:** Leiye Yu, Licong He, Bing Gan, Rujuan Ti, Qingjie Xiao, Hongli Hu, Lizhe Zhu, Sheng Wang, Ruobing Ren

## Abstract

As a critical sphingolipid metabolite, sphingosine-1-phosphate (S1P) plays an essential role in immune and vascular systems. There are five S1P receptors, designated as S1PR1-5, encoded in the human genome, and their activities are governed by endogenous S1P, lipid-like S1P mimics, or non-lipid-like therapeutic molecules. Among S1PRs, S1PR1 stands out due to its non-redundant functions, such as the egress of T and B cells from the thymus and secondary lymphoid tissues, making it a potential therapeutic target. However, the structural basis of S1PR1 activation and regulation by various agonists remains unclear. Here we reported four atomic resolution cryo-EM structures of Gi-coupled human S1PR1 complexes: bound to endogenous agonist d18:1 S1P, benchmark lipid-like S1P mimic phosphorylated Fingolimod ((S)-FTY720-P), or non-lipid-like therapeutic molecule CBP-307 in two binding modes. Our results revealed the similarities and differences of activation of S1PR1 through distinct ligands binding to the amphiphilic orthosteric pocket. We also proposed a two-step “shallow to deep” transition process of CBP-307 for S1PR1 activation. Both binding modes of CBP-307 could activate S1PR1, but from shallow to deep transition may trigger the rotation of the N-terminal helix of G_αi_ and further stabilize the complex by increasing the G_αi_ interaction with the cell membrane. We combine with extensive biochemical analysis and molecular dynamic simulations to suggest key steps of S1P binding and receptor activation. The above results decipher the common feature of the S1PR1 agonist recognition and activation mechanism and will firmly promote the development of therapeutics targeting S1P receptors.

## Introduction

Sphingolipids, named after the sphinx in Egypt to represent its mysterious role, include S1P, sphingosine, ceramide, and other complex sphingolipids like glycosphingolipids and sphingomyelins (1–3). S1P, acting as a bioactive lipid mediator, is mainly derived from the deacylation of ceramide or interconverted with sphingosine and secreted by vascular endothelial cells to the circulation predominantly (4–7). Intracellular S1P promotes cellular proliferation (8) and close links to a myriad of essential cellular processes, including immune cell trafficking (9), angiogenesis (10, 11), and vascular maturation (12). Plasma S1P helps maintain vascular integrity and regulate vascular leaks (13). Besides, S1P is identified as an early risk factor of lung cancer (14) and a crucial mediator of cardio-protection (15, 16). Thus, abnormal S1P production leads to the occurrence and progression of numerous severe diseases, such as metabolic syndrome, cancers, autoimmune disorders such as multiple sclerosis, and kidney and cardiovascular diseases (7, 17–20). Currently, the therapeutic molecules targeting S1PRs can be divided into two classes: the lipid-like S1P mimic such as (S)-FTY720-P (21) or the non-lipid-like molecules such as clinical drugs BAF-312 (Siponimod) (22), RPC-1063 (Ozanimod) (23), and CBP-307, which is still on clinical trials (Fig. 1A). (S)-FTY720-P is the *in vivo* phosphorylated product of Fingolimod to treat relapsing multiple sclerosis (MS) (3, 24). Siponimod and Ozanimod are also lunched to treat relapsing multiple sclerosis or ulcerative colitis in recent years (25, 26). Despite the broad indications and urgent need, the development of therapeutics is primarily limited by the high sequence similarity of S1PRs and less characterized functions of S1PR2-5. The inactive structure of S1PR1 bound with an antagonist was reported in 2012 (27). Recently, the crystal structure of active S1PR3 bound to Fab and endogenous agonist S1P and active state structures of S1PR1,3,5 bound to G_i_ complex and different agonists were reported (28–30). Here we reported four atomic resolution cryo-EM structures of G_i_-coupled human S1PR1 complexes bound to d18:1 S1P, (S)-FTY720-P, or CBP-307 in two binding modes. This structural information further supplements the binding details of different agonists and explores the activation mechanism of S1PR1, which will firmly promote the development of therapeutics targeting S1P receptors.

**Figure 1.**
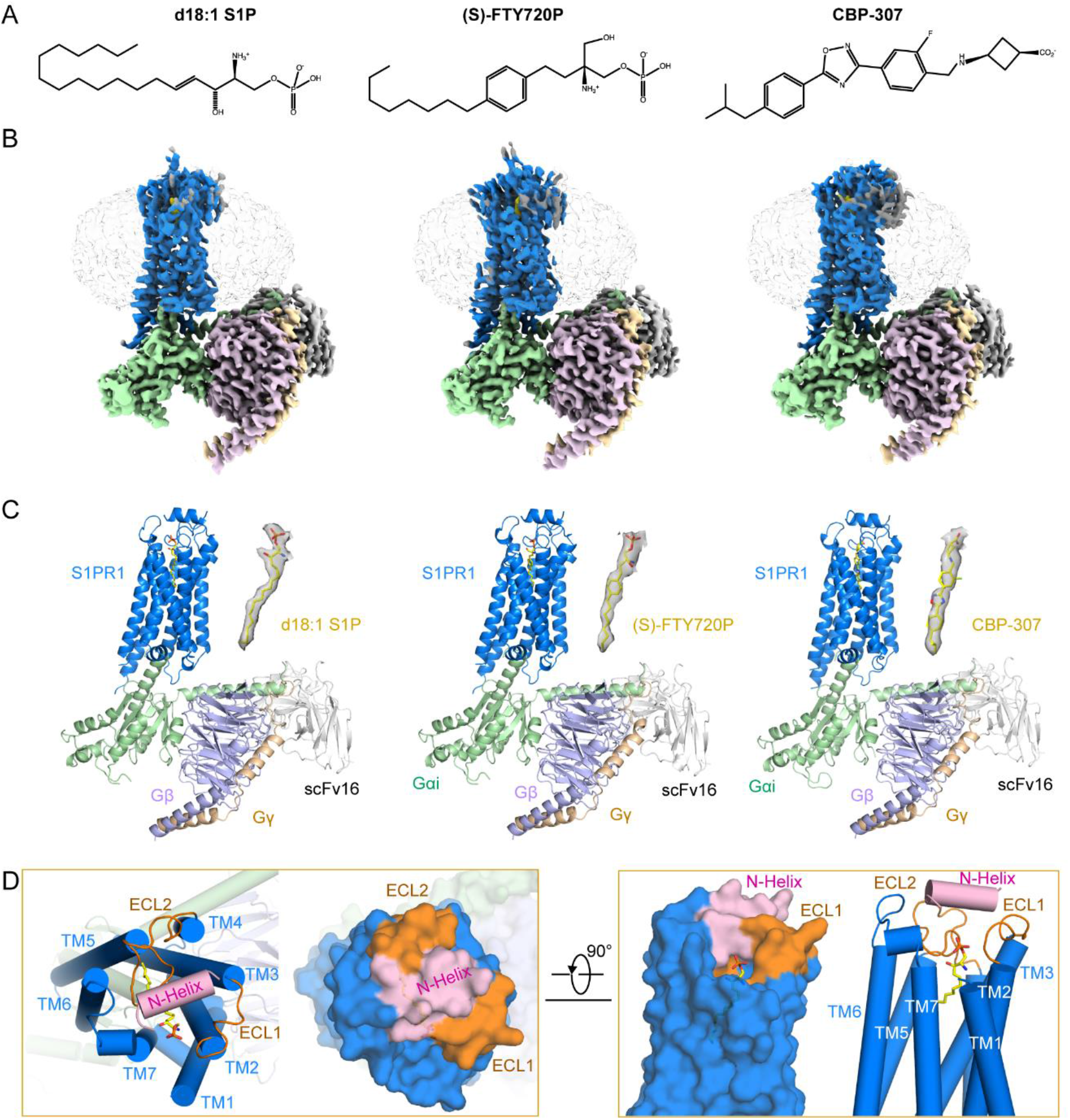
Cryo-EM maps and structures of human S1PR1 with Gi and agonist d18:1 1P, (S)-FTY720-P or CBP-307. **A**. Structural formulas of three agonists: d18:1 S1P, (S)-FTY720-P, and CBP-307. **B**. Cryo-EM maps of human S1PR1 with Gi and agonist d18:1 S1P, (S)-FTY720-P or CBP-307. The densities of S1PR1, G_αi_, G_β_, G_γ_, and scFv16 are shown in marine, pale green, pink, wheat, and grey, respectively. **C**. Structural models of human S1PR1 with Gi, scFv16, and agonist d18:1 S1P (PDB: 7VIE), (S)-FTY720-P (PDB: 7VIF) or CBP-307 (PDB: 7VIG). S1PR1, G_αi_, G_β_, G_γ_, and scFv16 are shown in marine, pale green, light blue, wheat, and grey. Ligands in density (yellow) are shown on the right side of the models. **D**. The extracellular lid of S1PR1 is shown in the top and side views (Surface and cartoon modes). N-Helix, ECL1/2, and TM helices are pink, orange, and marine, respectively.

## Results

### S1PR1-Gi complexes assembling and overall structures

For structure determination purposes, we engineered the human S1PR1 with an N-terminal BRIL fusion and the addition of affinity tags (an N-terminal FLAG tag and a C-terminal 10х His tag) (*SI Appendix*, Fig. S1A). S1PR1, G_αi1_, G_β_, and G_γ2_ were co-expressed in *sf9* insect cells using baculovirus to form the S1PR1-G_i_ complex. Agonist d18:1 S1P, (S)-FTY720-P, or CBP307, and the antibody fragment scFv16 were added during purification to enable a stable complex formation. The cell membrane was solubilized in lauryl maltose neopentyl glycol (LMNG) and cholesteryl hemisuccinate and then purified by two steps using FLAG and nickel-affinity beads sequentially. The concentrated samples were applied size-exclusion chromatography to yield a monodisperse complex that contained all the components (*SI Appendix*, Fig. S1B-D).

The vitrified complexes were imaged using a Titan Krios cryo-electron microscope. Images were processed to yield a final map at an overall resolution of 2.86 Å for d18:1 S1P bound (PDB code: 7VIE), that of 2.83 Å for (S)-FTY720-P bound (PDB code: 7VIF), and that of 2.98 Å and 2.89 Å for CBP-307 bound shallow (PDB code: 7VIH) and deep (PDB code: 7VIG) binding modes, respectively (Fig. 1B, *SI Appendix*, Fig. S2-5). The densities for the N terminus and transmembrane helices of receptor and G protein complexes were unambiguous determined, based on the well-traced α-helices and aromatic side chains (*SI Appendix*, Fig. S2-5). Due to the flexibility, the N-terminal BRIL fusion, part of the extracellular loops, and long C-terminal residues are invisible. It is worth mentioning that the densities of agonists are also well-defined (Fig. 1C).

S1PR1, with different agonists bound at the extracellular side and G_i_ complex bound at the intracellular side, demonstrate an active state super-complex, which is reminiscent of the structures of other G_i_-coupled class A GPCRs (29, 31). The structural analysis reflects some common features for other lipid receptors, such as cannabinoid receptors (32) and CRTH2 (33). The extracellular loops (ECL1-2) and the N-terminal “cap” region are coordinated to exclude agonists from the extracellular solvent (Fig. 1D, left). Besides, there is a cleft between TM1 and TM7 facing the membrane bilayer (Fig. 1D, right). These observations suggest a common mechanism that these lipidic molecules, such as S1P, cannabinoid, and prostaglandins, may first integrate into the lipid bilayer before binding to the receptors.

### Structures of d18:1 S1P and (S)-FTY720-P binding modes

Due to the structure similarity, d18:1 S1P and (S)-FTY720-P demonstrate almost identical fully extended conformations and form equivalent contacts in the orthosteric binding pocket of S1PR1 (Fig. 2A and B). The orthosteric pocket of S1PR1, which is highly conserved among all S1PRs (Fig. 2A, left, *SI Appendix*, Fig. S6), is divided by a polar region on the top (composed by N-terminal cap, ECL1 and 2) and a deep hydrophobic cavity (composed by TM3 and 5-7). These characters fit the zwitterionic nature of d18:1 S1P and (S)-FTY720-P well (Fig. 2A and B). Residues K34 and G106 form hydrogen bonds with the phosphate group of d18:1 S1P directly, while N101^2.60^ and S105 coordinate the amino group of d18:1 S1P (Fig. 2C). In addition, Y29 and R120^3.28^ may interact with the polar group of d18:1 S1P by van der Waals interactions, electrostatic interaction or possible indirect interactions such as water-mediated hydrogen bonds (*SI Appendix*, Fig. S7A). The interactions between the polar region of (S)-FTY720-P and S1PR1 also involve residues mentioned above. The three slight differences are: 1) Y29 but not K34 forms a hydrogen bond with the phosphate group of (S)-FTY720-P; 2) N101^2.60^ interacts with the hydroxymethyl group of (S)-FTY720-P, which does not exist in d18:1 S1P; 3) S105 does not interact with (S)-FTY720-P directly (Fig. 2C, *SI Appendix*, Fig. S7B). Notably, the key residue E121^3.29^, which interacts with ML056 in the S1PR1 inactive structure, also forms a hydrogen bond with the amino group of (S)-FTY720-P (Fig. 2D). In contrast, E121^3.29^ does not directly interact with the polar head of d18:1 S1P. It may interact with d18:1 S1P by van der Waals interactions and possible indirect interactions such as water-mediated hydrogen bonds, or stabilize the local conformation of the binding pocket through hydrogen bond network with surrounding residues (*SI Appendix*, Fig. S7A). To demonstrate the importance of these residues, we conducted the G_i_ dissociation assay for S1PR1 mutants. The results clearly showed that N101^2.60^A, R120^3.28^A, and E121^3.29^A aborted G_i_ coupling significantly for both d18:1 S1P and (S)-FTY720-P (Fig.2 E and F). Y29A and K34A compromised the potency of d18:1 S1P by more than 30 folds, whereas two folds that of (S)-FTY720-P compared to wild type S1PR1 (Fig.2 E and F, *SI Appendix*, Table 2). Our results suggested that hydrophilic residues on top of the pocket of S1PR1 play an essential role in stabilizing the polar head of d18:1 S1P and activating downstream G_i_ signaling.

**Figure 2.**
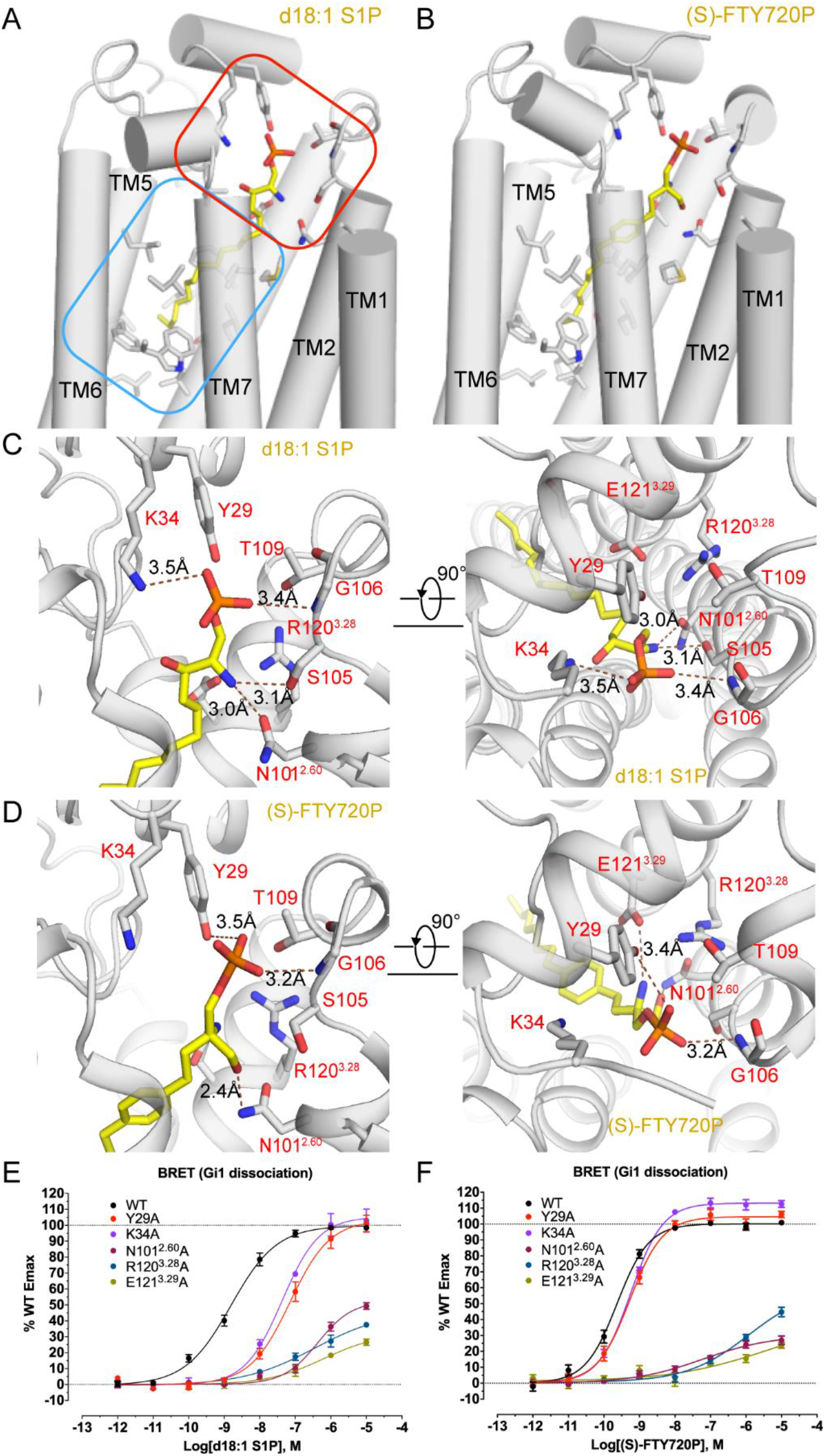
Orthosteric binding pockets of d18:1 S1P and (S)-FTY720-P in S1PR1. **A**. The binding pocket of d18:1 S1P in S1PR1. d18:1 S1P is shown in yellow. S1PR1 is gray with residues surrounding the binding pocket shown in sticks. The head and tail parts of the binding pocket are indicated by red and blue rectangular frames, respectively. **B**. The similar binding pocket of (S)-FTY720-P in S1PR1 compared to d18:1 S1P. (S)-FTY720-P and S1PR1 are shown the same color as in panel A. **C**. The key residues, including K34, N101^2.60^, S105, and G106, interact with the head group of d18:1 S1P. Side and top views are shown. The polar interactions are shown with dashed lines and indicated distances. **D**. The key residues, including Y29, N101^2.60^, G106, and E121^3.29^, interact with the head group of (S)-FTY720-P, which are shown the same color as in panel C. **E**. The effects of mutants Y29A, K34A, N101^2.60^A, R120^3.28^A, and E121^3.29^A of S1PR1 on d18:1 S1P induced Gi signal activation measured by Gi dissociation assay (BERT assay). All data are mean ± SEM of three independent experiments for wild-type or mutants. **F**. The effects of mutants Y29A, K34A, N101^2.60^A, R120^3.28^A, and E121^3.29^A of S1PR1 on (S)-FTY720-P induced Gi signal activation measured by Gi dissociation assay (BERT assay). All data are mean ± SEM of three independent experiments for wild-type or mutants.

The acyl chains of d18:1 S1P and (S)-FTY720-P are surrounded by numbers of hydrophobic residues, such as M124^3.32^, F125^3.33^, L128^3.36^, S129^3.37^, L195, F210^5.47^, W269^6.48^, L272^6.51^, L276^6.55^, and L297^7.39^ on TM3 and TM5-7 (*SI Appendix*, Fig. S7C). The superposition of d18:1 S1P and (S)-FTY720-P bound S1PR1 structures showed that acyl chains of d18:1 S1P extend deeper than the tail of (S)-FTY720-P (*SI Appendix*, Fig. S7C). We also examined the effects of these hydrophobic residues on agonist binding and G_i_ activation. Unsurprisingly, the potency of d18:1 S1P and (S)-FTY720-P to most of the S1PR1 mutants are decreased (*SI Appendix*, Table 2). W269^6.48^, L272^6.51^A, L276^6.55^A, and L297^7.39^A showed the most significant changes (*SI Appendix*, Fig. S7D, and E). Together, these interactions make a high binding affinity for d18:1 S1P and (S)-FTY720-P to S1PR1. It’s worth mentioning that the signature residues for GPCR activation, F210^5.47,^ and W269^6.48^, are located on the bottom of the pocket, indicating the critical role of the S1P acyl chain for S1PR1 activation.

### Activation mechanism of S1PRs

We firmly believe that, together with the antagonist ML056 bound inactive S1PR1 structure (27) and the active S1PR1 structures bound to G_i_ and various agonists, it is a proper prototype to propose the activation mechanism S1P receptors. Here we used d18:1 S1P bound structure as the representative active state to analyze the conformational changes during the S1PR1 activation process. In the superposition of active and inactive S1P1R structures, the overall r.m.s.d is 1.222 Å over 269 residues majorly located on the TM region. The extracellular half of S1PR1 showed an unapparent change that the N-terminus of TM1 moved towards TM7 by 2.7 Å (C_α_ of Val50 as reference). In contrast, the conformation of intracellular half significantly changed with TM6 moving away from TM7 by 8.3 Å (C_α_ of Lys250 as reference), and TM7 swinging inward by 3.4 Å (C_α_ of Leu313 as reference), resulting in the G_αi_ coupling. (Fig. 3A).

**Figure 3.**
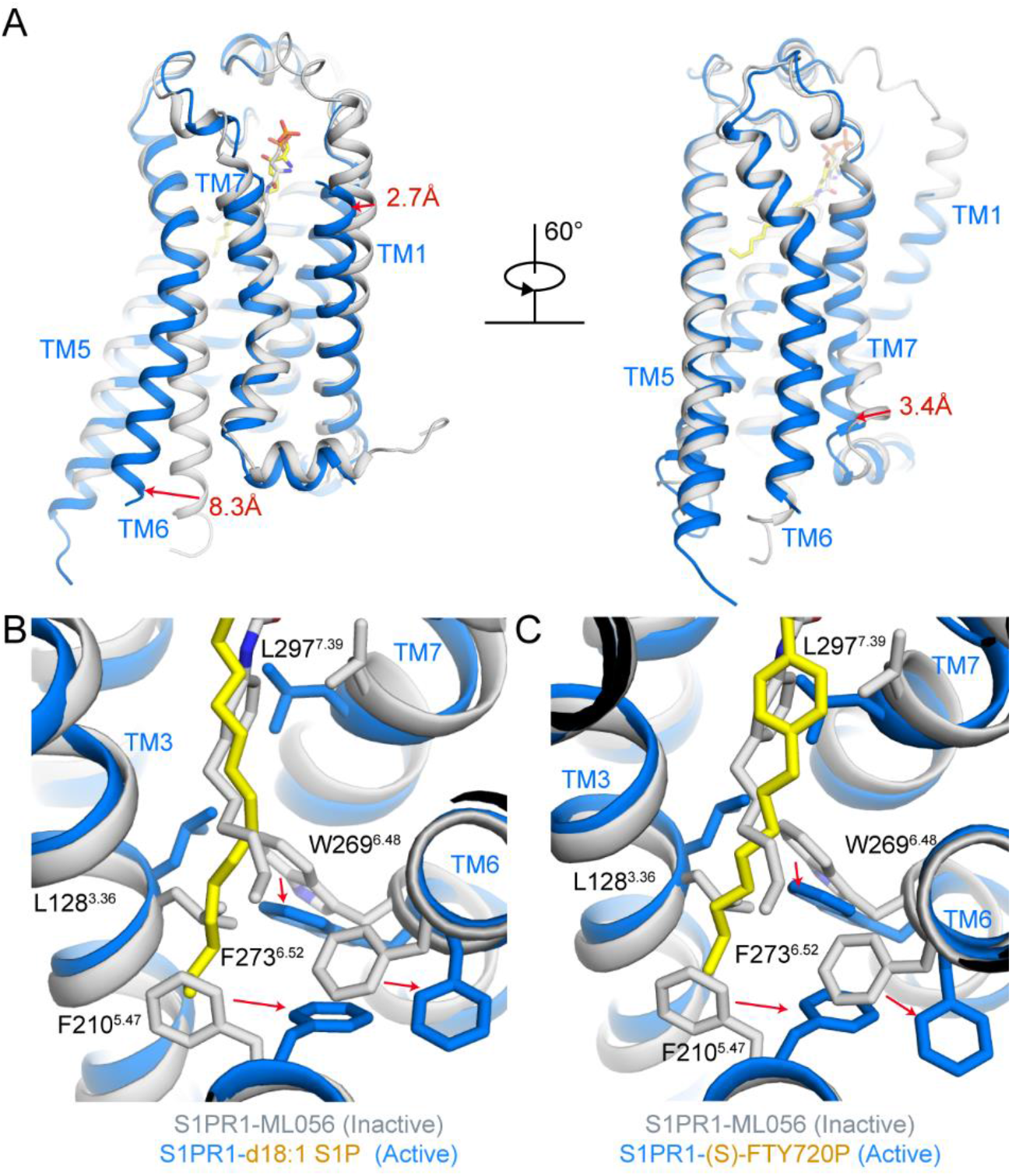
Conformational changes between active and inactive forms of S1PR1. **A**. TM helix shifts between S1PR1 (colored in marine) in the active form of S1PR1-Gi-d18:1 S1P complex (PDB: 7VIE) and S1PR1 (colored in gray) in inactive form of S1PR1-ML056 complex (PDB: 3V2Y). Red arrows indicate significant shifts of TM1, TM6, and TM7 with distances. **B**. Conformational changes of key residues at the bottom of the pockets between active (d18:1 S1P bound) and inactive forms are shown in marine and white. d18:1 S1P and ML056 are shown in yellow and white, respectively. Red arrows indicate the directions of conformation changes of L128^3.36^, F210^5.47^, W269^6.48^, F273^6.52^, and L297^7.39^ are shown in sticks. The directions of conformation changes of F210^5.47^, W269^6.48^, and F273^6.52^. **C**. Conformational changes of key residues at the bottom of the pocket between active ((S)-FTY720-P bound) and inactive forms are shown in marine and white. (S)-FTY720-P and ML056 are shown in yellow and white, respectively. Residues of L128^3.36^, F210^5.47^, W269^6.48^, F273^6.52^, and L297^7.39^ are shown in sticks. Red arrows indicate the conformational changes of F210^5.47^, W269^6.48^, and F273^6.52^.

Further careful examines of signature residues in the orthosteric pocket of S1PR1 showed a well-organized spatial rearrangement of a few side chains. The most attractive residue is F210^5.47^, located at the bottom of the pocket. In the inactive structure, F210^5.47^ points to TM3 but does not interact with ML056. However, upon d18:1 S1P binding, F210^5.47^ rotates about 130°to avoid a stereo clash with the acyl chain of d18:1 S1P inserted deeper than ML056. Then, the F210^5.47^ rotation results in the rotation of F273^6.52^ and pushes TM6 moving outward. L128^3.36^ also rotates away from W269^6.48^ to adapt the accommodation of d 18:1 S1P and weaken the interactions between TM3 and TM6. It will facilitate the activation of S1PR1 (Fig. 3B). A similar observation is in the (S)-FTY720-P bound structure (Fig. 3C). Thus, the initiation of S1PR1 activation is triggered most likely by touching L128^3.36^ and F210^5.47^ upon agonist binding. The slight conformational change of the extracellular half of TM7 is due to the movement of L297^7.39^ to form hydrophobic interaction with the acyl chain of d18:1 S1P and (S)-FTY720-P (Fig. 3B and C).

### Structures of CBP-307 binding modes

CBP-307 is a synthetic non-lipid-like molecule targeting S1PR1/4/5 specifically. In the process of structure determination, there are two classes of particles with comparable map quality showing slight differences by hetero refinement (*SI Appendix*, Fig. S4A). After carefully checking the densities of the receptor, G_i_, and CBP-307, we found that CBP-307 adopted two distinct binding modes (Fig. 4A). Then, two structural models were built and superimposed to analyze the structural differences. Only the receptor was used for alignment, and the overall r.m.s.d is 0.313 Å over 257 residues, indicating the identical conformation of S1PR1 in these two binding modes. We named “shallow” and “deep” to define these two binding modes because CBP-307 in the deep mode is inserted 1.8 Å deeper into the pocket than that in the shallow mode (Fig. 4B). Densities of CBP-307 in these two structures were shown (Fig. 4C).

**Figure 4.**
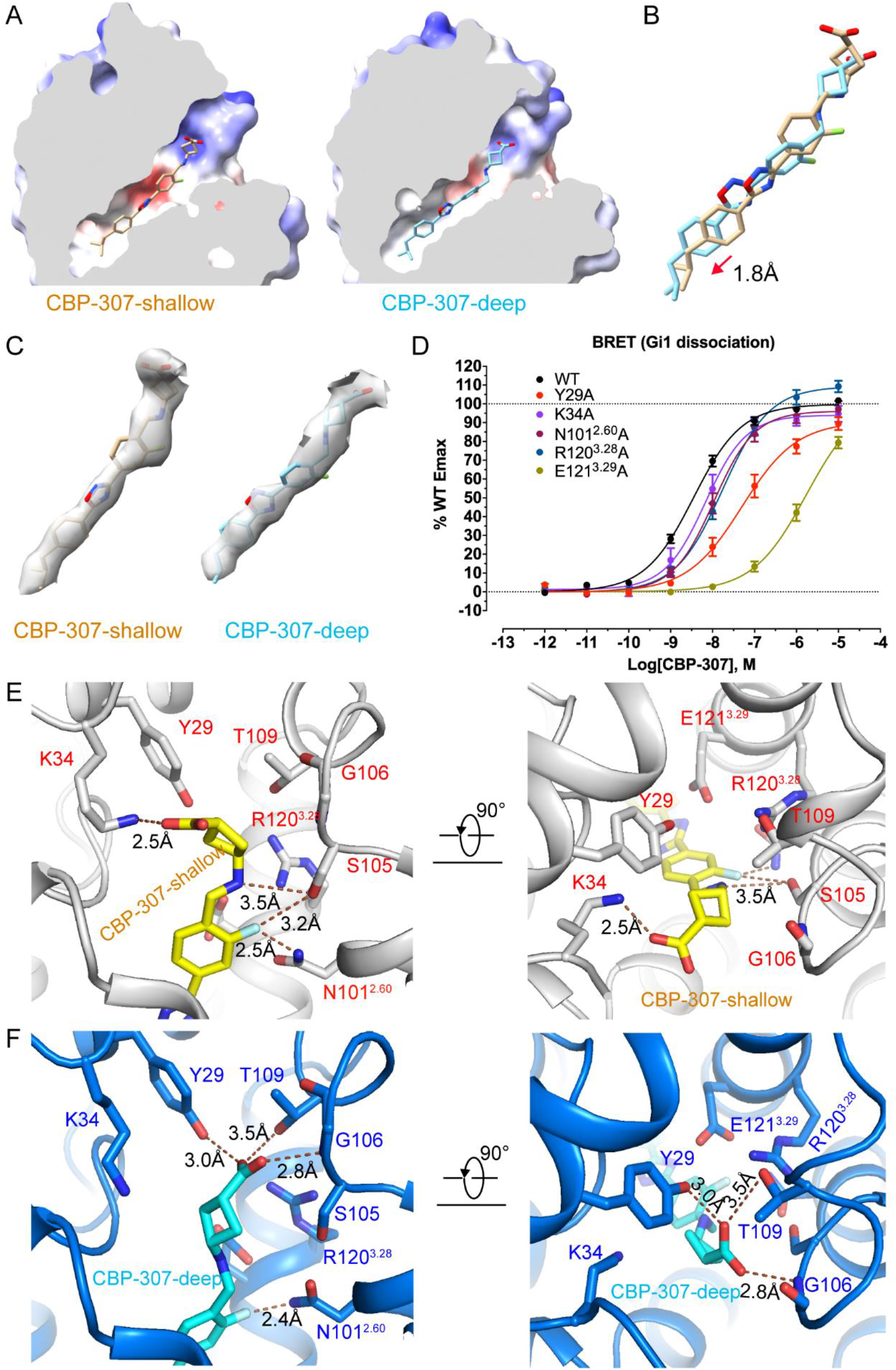
CBP-307 binding with S1PR1 in two binding modes. **A**. A cutaway representation of two binding modes of CBP-307 bound S1PR1 at the level of binding pocket, and colored according to surface electrostatic potential with cut surface shown in gray. CBP-307 in shallow mode is colored in wheat. CBP-307 in deep mode is colored in cyan. CBP-307 is shown in sticks. **B**. Superposition of two binding modes of CBP-307. CBP-307 in deep mode inserts into the pocket deeper with a 1.8 Å distance than shallow mode. CBP-307 is shown and colored as A. **C**. CBP-307 is modeled in two density maps. **D**. The effects of Y29A, K34A, N101^2.60^A, R120^3.28^A, and E121^3.29^A mutations of S1PR1 on CBP-307 induced Gi signal activation measured by Gi dissociation assay (BERT assay). All data are mean ± SEM of three independent experiments for wild-type or mutants. **E**. The key residues interacted with the polar groups of CBP-307 in shallow mode. S1PR1 and CBP-307 are shown in gray and yellow, respectively. CBP307, Y29, K34, N101^2.60^, S105, G106, T109, R120^3.28A^, and E121^3.29^ are shown in sticks. Dashed lines indicate the polar interactions between CBP-307 and K34, N101^2.60^, and S105 with distances. **F**. The key residues interacted with the polar groups of CBP-307 in deep mode. S1PR1 and CBP-307 are shown in marine and cyan, respectively. CBP307, Y29, K34, N101^2.60^, S105, G106, T109, R120^3.28A^, and E121^3.29^ are shown in sticks. Dashed lines with distances indicate the polar interactions between CBP-307 and Y29, N101^2.60^, G106, and T109.

Similar to d18:1 S1P and (S)-FTY720-P structures, residues Y29, K34, N101^2.60^, G106, R120^3.28^, and E121^3.29^ forms a hydrophilic pocket to coordinate the carboxyl group of CBP-307. The functional mutagenesis data also suggested the importance of these polar residues (Fig. 4D, *SI Appendix*, Table 2). For hydrophobic residues, the G_i_ dissociation assay showed that W269^6.48^A, L272^6.51^A, and L276^6.55^A weakened CBP-307 mediated G_i_ signal similar to d18:1 S1P (*SI Appendix*, Fig. S8A and B, and Table 2). Notably, F125A also weakened the CBP-307 mediated G_i_ signal significantly, indicating its potential role in interacting with the benzene ring of CBP-307 (*SI Appendix*, Fig. S8A and B, and Table 2).

After carefully comparing the detailed interactions between CBP-307 and S1PR1 in two binding modes, we noticed that the surrounding residues that interacted with CBP-307 in two structures were significantly different. In the shallow mode, the carboxyl group of CBP-307 forms a hydrogen bond to K34, and the fluorine interacts with N101^2.60^ and S105 due to its negatively charged property (Fig. 4E). Besides, the amino group forms a hydrogen bond with S105 (Fig. 4E). In the deep mode, the carboxyl group of CBP-307 forms extensive hydrogen bonds to Y29, G106, and T109 (Fig. 4F). Additionally, the deeper insertion of CBP-307 makes its amino group closer to E121^3.29^ to form stronger interaction than the shallow mode (*SI Appendix*, Fig. S9A and B). All observations suggest that, in the deep mode, CBP-307 binds to the hydrophilic part of the S1PR1 pocket stronger and similar to the binding modes of d18:1 S1P and (S)-FTY720-P to the receptor (*SI Appendix*, Fig. S9A and B). Extensive molecular dynamic (MD) simulations were also performed to confirm our speculation further. The result showed that the CBP-307 in the shallow mode inserted deeper after about 50ns and stayed in a similar position to the deep mode for longer than 1100ns, consistent with our observations in structures (*SI Appendix*, Fig. S10A and B).

### Activation of S1PR1 by CBP-307

In the superposition of d18:1 S1P bound and two CBP-307 bound structures, the isobutyl group of CBP-307 in the deep binding mode occupies the space of the acyl end of d18:1 S1P, acting as the similar function in S1PR1 activation (Fig. 5A). Notably, the isobutyl group of CBP-307 in the shallow mode has less stereo clash with F210^5.47^ than in the deep mode, although the receptor is still activated (Fig. 5B). In both binding modes, the interfaces between S1PR1 and the α5 helix of G_αi_ are almost identical. The residues R78^2.37^, R142^3.50^, and K250^6.29^ on S1PR1 form hydrogen bonds with the carbonyl oxygen of K348, D345, C351, and the carboxyl group of D341 on G_αi_, respectively. The side chains of N153 on ICL2 of S1PR1 and D350 on G_αi_ forms a hydrogen bond. The carbonyl oxygen of M149 on ICL2 of S1PR1 interacts with N347 on G_αi_ (Fig. 5C). This extensive hydrogen bond network forces G_αi_ to bind to S1PR1 tightly.

**Figure 5.**
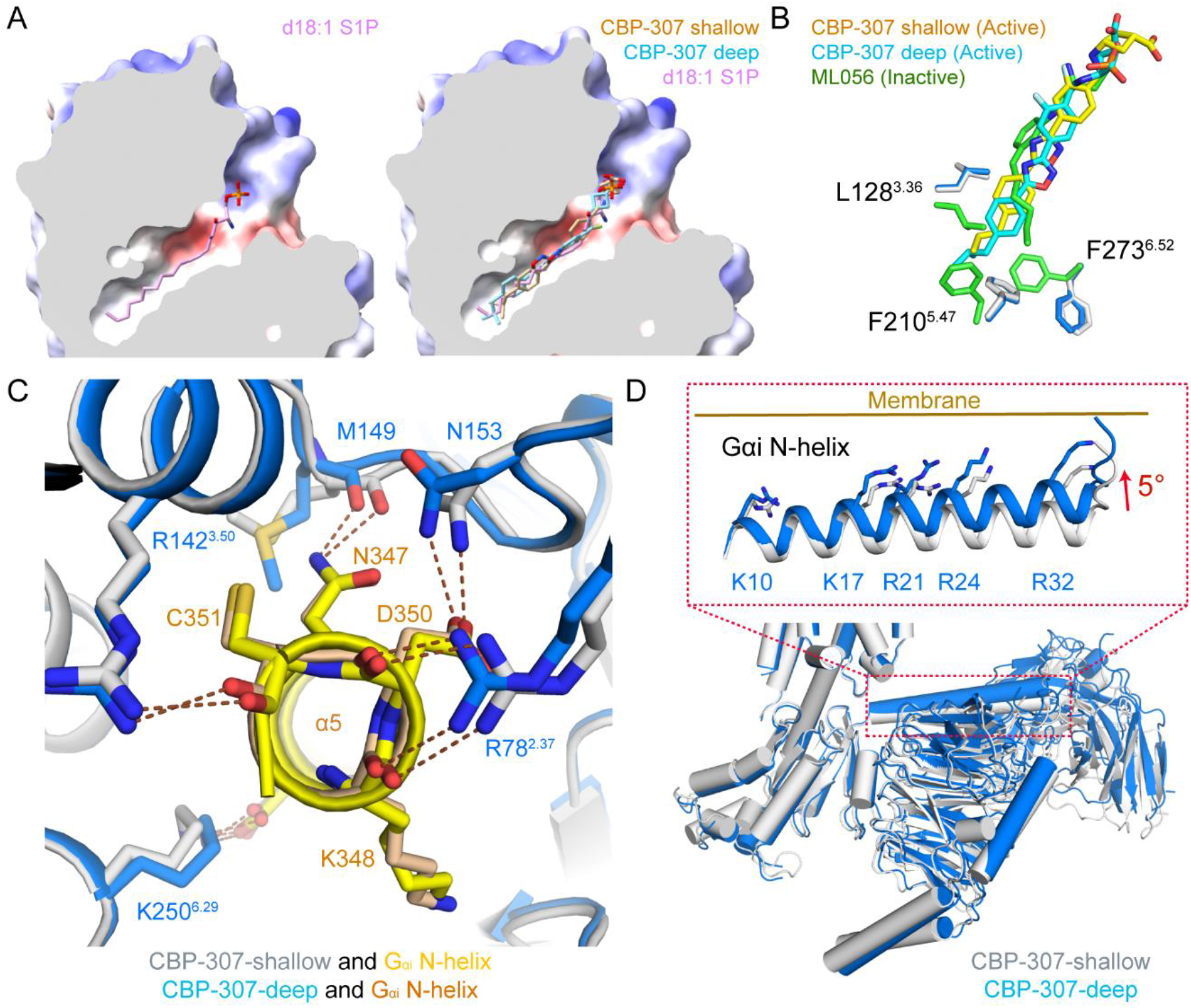
Activation of S1PR1 by CBP-307. **A**. A cutaway representation of d18:1 S1P in S1PR1-G_i_-d18:1 S1P model and superimposed two CBP-307 binding modes with d18:1 S1P in S1PR1-G_i_-d18:1 S1P model. The cutaway surface of S1PR1 with d18:1 S1P is shown at the level of binding pocket, and colored according to surface electrostatic potential with cut surface shown in gray. d18:1 S1P and CBP-307 are shown in sticks. d18:1 S1P, CBP-307 (shallow), and CBP-307 (deep) are shown in violet, wheat, and cyan, respectively. **B**. Superposition of ML056 with CBP-307 inactive and two CBP-307 bound active forms. ML056, CBP-307-shallow, and CBP-307-deep are shown in sticks colored in green, yellow, and cyan, respectively. Residues near the isobutyl group, including L128^3.36^, F210^5.47^, and F273^6.52^ in S1PR1-ML056, S1PR1-CBP-307(shallow), and S1PR1-CBP-307(deep), are shown in sticks and colored in green, gray, and marine, respectively. **C**. The polar interactions of Gi and S1PR1 with CBP-307 in two modes are similar. S1PR1-CBP-307 (shallow) and S1PR1-CBP-307 (deep) are shown in gray and marine, respectively. The helix 5 of G_αi_ in S1PR1-CBP-307 (shallow) and S1PR1-CBP-307 (deep) are shown in yellow and wheat, respectively. Residues on S1PR1 and helix 5 of G_αi_ are shown in sticks. **D**. The N-terminal helix of G_αi_ riched in positive charge residues at a closer position to the membrane in the CBP-307 deep mode structure. CBP-307 deep and shallow modes are colored in marine and gray, respectively. The positive charge residues (K10, K17, R21, R24, and R32) on the N-terminal helix of G_αi_ are shown in stick in an inset.

However, the N-terminal helices of G_αi_ in these two structures have a 5° rotation (Fig. 5D, *SI Appendix*, Fig. S11A). This rotation leads to the following movement of G_β_ and G_γ_. The N-terminal helix of G_αi_ is rich in positively charged residues such as K10, K17, R21, R24, and R32 (Fig. 5D). These positive charge residues pointed in the same direction interact with membrane lipids. The N-terminal helix of G_αi_ in the deep model is closer to the membrane and may help to stabilize the S1PR1-G_i_ complex. Structure alignment also suggested that the overall structure of the CBP-307 deep mode is similar to the structure of the d18:1 S1P bound S1PR1 complex (*SI Appendix*, Fig. S11B) and the four structures reported by *Shao* group (*SI Appendix*, Fig. S11C)(29). With the analysis of CBP-307 binding modes and the conformational change of G protein complexes, we propose a two-step model of S1PR1 activation by CBP-307. Two structures represented two relatively stable conformations during S1PR1 activation. Firstly, CBP-307 binds to the shallow binding site, activates S1PR1, and recruits G_i_ to form a complex. Then CBP-307 moves deeper to trigger the G_i_ complex rotation towards the membrane and further stabilizes the S1PR1-G_i_ complex.

## Discussion

Although a few structures of inactive S1PR1, active structures of S1PR1, S1PR3, and S1PR5 were reported, and the selectivity of alkyl length of S1P analogs are carefully examined (34, 35), it is still valuable to determine the S1P bound S1PR1 active structure. Moreover, different agonists’ recognition and activation of S1PRs need to be studied extensively. Here we reported G_i_-coupled active structures of S1PR1 with three distinct agonists, revealing the agonist recognition and receptor activation mechanism of S1PR1. The previous MD simulation studies suggested that lipidic agonists may enter the pocket through the cleft between the extracellular half of TM1 and TM7 (36). Residue E121^3.29^ may guide S1P penetration and stabilization, while W269^6.48^ is moving towards TM7 results in receptor activation through forming a hydrogen bond network in the intracellular half of TM7 (37). The space between TM3 and TM6 is increased upon G protein complex binding. All these speculations are also observed in the active state structures we reported. Since key residues we identified with composing the orthosteric binding pockets and mediating receptor activation, including Y29, K34, N101^2.60^, S105, G106, R120^3.28^, E121^3.29^, F125^3.33^, F210^5.47^, W269^6.48^, L272^6.51^ (L259^6.51^ in S1PR2), L276^6.55^ (F263^6.51^ in S1PR3), and L297^7.39^ (F274^7.39^ in S1PR2), are almost identical among S1PRs, they would share the very similar activation mechanism upon d18:1 S1P binding.

CBP-307 is currently being evaluated in a global Phase 2 clinical study in moderate to severe ulcerative colitis and Crohn’s disease (38). As reported, CBP-307 is a specific agonist for S1PR1/4/5 and has significantly higher potency for S1PR1 than S1PR4 and S1PR5, which is also confirmed by the G_i_ dissociation assay we performed (*SI Appendix*, Fig. S12A). The reduction of non-specific activation of S1PR2 will protect the vascular integrity, and that of S1PR3 may reduce the risk of fibrosis and bronchoconstriction. However, the detailed mechanism of CBP-307 selectivity is still poorly understood. We compared CBP-307 bound S1PR1-G_i_ structure (deep mode) to Siponimod bound S1PR5-G_i_ and the d18:1 S1P bound S1PR3-G_i_ structures reported by *Shao* group (29, 30). Residues composed of the orthosteric binding pocket of S1PR1 and S1PR5 are highly conserved (*SI Appendix*, Fig. S6). Only a few polar residues adopted different rotamers to coordinate the polar head of different agonists (*SI Appendix*, Fig. S12B). It could explain the cross-activity of CBP-307 to S1PR1 and S1PR5 to some extent. The corresponding residue of L276^6.55^ in S1PR1 is F263^6.51^ in S1PR3. Another two residues in S1PR1, S129^3.37^and V132^3.40^ (G123 and T126 in S1PR3), are variable (*SI Appendix*, Fig. S12C). Considering the importance of L276^6.55^ in S1PR1 activation and the locations of S129^3.37^and V132^3.40^, further investigation combining structural information, MD simulations, and pharmacological analysis is needed. When we examined the orthosteric binding pocket of S1PR1 to different agonists, two unoccupied sub-pockets were observed in d18:1 S1P bound structure (*SI Appendix*, Fig. S12D). These two sub-pockets are different from those identified in S1PR3 (30). More strikingly, the second sub-pocket is also empty in the other three S1PR1 structures (*SI Appendix*, Fig. S12E). This pocket can be occupied by agonists to improve the potency and selectivity by designing derivatives of S1PR1 agonists. Collectively, since the structure-based drug design has been applied to many GPCRs, it is vital to obtain structures of S1P receptors in multiple conformations bound with diverse ligands to benefit drug discovery targeting S1PRs by reducing side effects or extending indications.

## Materials and Methods

Detailed experimental procedures used in this study, including cloning, cell culture, recombinant protein expression and purification, cryo-EM data collection, and structure determination, are available as *SI Appendix, Experimental Procedures*. All materials and detailed protocols are available on request from the corresponding author.

## Data Availability

All relevant data are available from the authors and/or included in the manuscript. Atomic coordinates and EM density maps of four structures have been deposited in the Protein Data Bank and the Electron Microscopy Data Bank, respectively. The PDB and EMDB codes are available in *SI Appendix, Table S1*. The structure models are presented by Pymol (https://pymol.org/2/), and the density maps are presented by Chimera and Chimera X (39, 40).

## Author Contributions

R. R conceived the project and designed all experiments. L. Y., B. G., L. H., R. T., and Q. X. performed all experiments. B. G. purified receptor-G protein complex samples. L. Y. prepared the Cryo-EM grids, collected the EM data, and determined structures. L. H. performed pharmacology assays and analyzed the data. Q. X. and H. H. helped to validate structural models. R. T. performed the dynamic simulation. S. W. supervised the pharmacology assays. L. Z. supervised the dynamic simulation. All authors analyzed the data and contributed to manuscript preparation. R. R. and L. Y. wrote the manuscript.

## Acknowledgments

We are grateful to Connect Biopharm for donating agonist CBP-307 and critical discussion. We also thank Kobilka cryo-EM Center at the Chinese University of Hong Kong, Shenzhen, for supporting EM data collection. This work was supported by funds from the National Natural Science Foundation of China (Project 31971218 to R. R, 32071197 to S. W., and 31971179 to L. Z.), Ministry of Science and Technology of China (2020YFA0509102 to S. W.), the Science, Technology, and Innovation Commission of Shenzhen Municipality (Projects JCYJ-20180307-151618765 and JCYJ-20180508-163206306 to R. R, JCYJ-2020010915-0003938, and RCYX20200714114645019 to L. Z.,), Shanghai Science and Technology Committee (19ZR1466200 to S. W.), and the Thousand Talents Plan-Youth and Shanghai Rising-Star Program (20QA1410600 to S. W.). R. R. and L. Z. were also supported in part by Presidential Fellowship and B. Gan was supported by Ganghong Youth Scholarship at the Chinese University of Hong Kong, Shenzhen.

