## Supplemental Materials for "Structural Insights into Sphingosine-1-phosphate Receptor Activation"

### Supplementary Information

#### Material and Methods

##### Compounds and reagents

CBP-307 was a generous gift of Connect Biopharma. d18:1 S1P (Sphingosine-1-Phosphate (d18:1)) and (S)-FTY720-P were purchased from Avanti Polar Lipids and Toronto Research Chemical, respectively. Other reagents were purchased from Sigma-Aldrich and Sangon Biotech.

##### Constructs

The full-length wild-type human S1PR1 (residues M1-S382) was cloned into a pFastBac vector (Invitrogen) with a HA signal peptide, an N-terminal Flag tag, and a C-terminal 10xHis tag. To increase the expression level of S1PR1, a BRIL protein was fused at the N-terminus of the receptor after the 3C protease site. There was no more modification in the S1PR1 sequence. The wild-type human $G_{\alpha i1}$ and a single-chain antibody scFv16 were cloned into pFastBac vectors. The wild-type human $G_{\beta 1}$ with an N-terminal 6×His-tag and $G_{\gamma 2}$ were cloned into a pFastBac-Dual vector. The construct was generated with a standard PCR-based strategy and homologous recombination (CloneExpress One Step Cloning Kit, Vazyme).

##### Expression of complex

The complex was expressed in *Spodoptera frugiperda* (Sf9) insect cells using the Bac-to-Bac system (Invitrogen). Baculovirus preparation was accomplished based on the baculovirus system manual (Invitrogen). For expression, the Sf9 cells were cultured in a serum-free medium (Sf-900$^{TM}$ II SFM (1x), Gibco). At a density of 2.0-2.5 million cells per milliliter, the Sf9 cells were co-infected with three different viruses (S1PR1 receptor, WT human $G_{\alpha i1}$, WT human $G_{\beta 1\gamma 2}$) at the ratio of 1:1:1. The infected cells were harvested by centrifugation (2,000g, 15 minutes, 4℃) after 48h. The cell pellets were frozen in liquid nitrogen and stored at −80 °C for use.

**Expression and purification of scFv16**

scFv16 were expressed from *Trichoplusia ni* (Hi5) insect cells as a secreted protein using the baculovirus infection system and the purification process as previously reported (1). The Hi5 insect cells were grown in serum-free medium (SIM HF) to a density of 3.0-4.0 million cells per ml and then infected with scFv16 virus produced in Sf9 cells at a ratio of 1:50 (virus volume vs. cells volume). The media expressing scFv16 was isolated from the cell pellets by centrifugation (2,000g, 15 minutes, 4℃) after 96h. The collected media was pH balanced to pH 8.0 byadding tris base powder. Chelating agents were quenched by adding 1 mM nickel chloride and 5 mM calcium chloride. The mixture was incubated at room temperature (25 °C) for 1h by stirring constantly. The resulting precipitates were removed by centrifugation (5251g, 30 minutes, 4℃), and the supernatant was incubated with Ni-sepharose resin (GE Healthcare) for 1h at room temperature by stirring constantly. The resin was collected by centrifugation (800g, 10 minutes, 4℃) after incubation and transferred to the gravity column for further procedures. The resin was then washed with the washing buffer (20 mM HEPES pH7.5, 100 mM NaCl, and 20 mM Imidazole). The scFv16 protein was eluted with the elution buffer (20 mM HEPES pH 7.5, 100 mM NaCl, and 250 mM Imidazole), and the carboxyl-terminal His-tag was cleaved by incubating with HRV-3C protease at 4℃ for 1h or more. The cleaved protein was further purified by size exclusion chromatography using a Superdex 200 Increase 10/300 GL column (GE healthcare) with running buffer 20 mM Hepes pH 7.5 and 100 mM NaCl. The targeted scFv16 peak fractions were pooled, concentrated, and flash-frozen in liquid nitrogen until use.

**Purification of complex**

The complex cell pellets were thawed and resuspended in 25 mM HEPES pH 7.5, 150 mM NaCl, 5% Glycerol, 10 mM $MgCl_2$, 20 mM KCl, 5mM $CaCl_2$ and 1 mM $MnCl_2$ supplemented with Protease Inhibitors (100 μM PMSF, 2 μg/mL Aprotinin, and 2 μg/mL Pepstatin). The suspension was homogenized by douncing ~30 times. Then, for the CBP307-$G_i$-ScFv16 complex, 10 μM CBP-307 and 25 mU/mL apyrase (NEB) was added to the suspension; for the d18:1 S1P-Gi-ScFv16 complex, 2 μM d18:1 S1P and 25 mU/mL apyrase (NEB) was added to the suspension; for the (S)-FTY720-P-Gi-ScFv16 complex, 2 μM FTY-720-P and 25 mU/mL apyrase (NEB) was added to the suspension. To get a stable complex, CBP-307, d18:1 S1P, or (S)-FTY720-P were added at a final concentration of 10 μM, 2 μM, and 2 μM all through the further purification procedure. The suspension with apyrase and ligand was incubated for 1h at room temperature. Then, 1% n-dodecyl-β-D-maltoside (DDM) (Bluepus) and 0.1% (w/v) cholesteryl hemisuccinate (CHS) (Anatrace) was added to solubilize the membrane. After incubating with the detergent for 2 h at 4 °C. The insoluble material was removed by centrifugation (39,191g, 30 minutes, 4°C). The supernatant was collected and incubated with Flag G1 affinity resin (GenScript) for 1h at 4 °C. The Flag resin was washed with washing buffer (25 mM HEPES pH 7.5, 150 mM NaCl, 5% Glycerol, 5 mM $MgCl_2$, 5mM $CaCl_2$, 0.01% (w/v) lauryl maltose neopentylglycol (LMNG) (Anatrace) and 0.001% (w/v) CHS (Anatrace)), then eluted with the same buffer plus 200 μg/mL Flag peptide (GenScript). The elution was loaded onto Ni-NTA resin (Qiagen) for further purification. The Ni-NTA resin was washed with washing buffer (25 mM HEPES pH 7.5, 150 mM NaCl, 5 mM $MgCl_2$, 25 mM Imidazole, 0.01% (w/v) lauryl maltose neopentylglycol (LMNG) and 0.001% (w/v) CHS), then the complex was eluted with elution buffer (25 mM HEPES pH 7.5, 150 mM NaCl, 5 mM $MgCl_2$, 250 mM Imidazole, 0.01% (w/v) lauryl maltose neopentylglycol (LMNG) and 0.001% (w/v) CHS). The final complex elution was concentrated to less than 2 mL using an Amicon Ultra Centrifugal Filter (MWCO 100 kDa) and then incubated with excess purified scFv16 for 2h on ice. The scFv16-bound complex was subjected to Superdex 200 Increase 10/300 GL column (GE healthcare) with running buffer (25 mM HEPES pH 7.5, 150 mM NaCl, 0.00075% (w/v) LMNG, 0.00025% (w/v)

GDN (Anatrace), 0.000075% (w/v) CHS, 100 μM TCEP) to remove uncoupled receptor and excess scFv16. The monomeric peak fractions containing receptor–$G_i$-scFv16 complex were pooled and concentrated to the final concentration of 10-13 mg/mL for cryo-EM grid preparation.

**Cryo-EM sample preparation and data acquisition**

3.5 μL of 12, 10.9, 13 mg/mL the concentrated S1PR1S (S1PR1-$G_i$- d18:1 S1P), S1PR1F (S1PR1-$G_i$-(S)-FTY720-P) and S1PR1C (S1PR1-$G_i$-CBP-307) complex was applied to glow-discharged holey carbon-coated grids (Quantifoil 300 mesh, Au R1.2/1.3). The grids were blotted for 3s and flash-frozen in liquid ethane using a Vitrobot (Mark IV, Thermo Fisher Scientific). Images were recorded on a 300kV Titan Krios G3i electron microscope (Thermo Fisher Scientific) equipped with Gatan K3 Summit direct detector and a GIF Quantum energy filter (slit width 20 eV). Movie stacks were collected using SerialEM (2) in counting mode at a magnification of 105,000x with the corresponding pixel size of 0.83Å. Movies stack with 50 frames of complex S1PR1S, S1PR1F, and S1PR1C were exposed for 2.5 s, 3 s, and 3 s. Movies of complex S1PR1S, S1PR1F, and S1PR1C were recorded at a dose rate of 15.47/17.36/18.32, 12.27, and 13.0 e/px/s separately, corresponding to a total dose of about 56.14/62.30/66.50 e/Å$^2$, 53.43 e/Å$^2$, 56.61 e/Å$^2$. The defocus range was set from −1.1 μm to −1.9 μm. A total of movie stacks of 5,997, 6,345, and 5,517 were collected for complex S1PR1S, S1PR1F and S1PR1C.

**Data Processing**

Movies frames were aligned using MotionCor2 (3) with 5 by 5 patches. Micrograph contrast transfer function (CTF) estimations were performed by CTFFind4 (4) using micrographs without dose-weighting. The dose-weighted micrographs were used for particle picking and further processing. Particles were automatically picked by Gautomatch (https://www.mrc-lmb.cam.ac.uk/kzhang/Gautomatch). For data of S1PR1C, S1PR1S and S1PR1F, 4,706,164 particles, 5,545,033 particles and 4,706,164 particles were picked. Micrographs with estimated Ctf Max Resolution (> 5 Å) and Ctf Figure of Merit (> 0.25) were selected for further processing.

For data of S1PR1S, due to dose increasing during data collection, movies alignment using MotionCor2 were done in three job with different dose per frame. Totally, 5,407 micrographs were selected with 5,007,355 particles. Particles were extracted using Relion3.1 (5) with a box size of 72 pixels and a pixel size of 3.32 Å. Iterative 2D classifications were performed using cryoSPARC v.2.14.2 (6). 2,125,571 particles were selected. One round ab initio reconstruction was performed. Particles of two good classes were re-centered and re-extracted separately by Relion with a box size of 288 pixels and a pixel size of 0.83 Å. Non-Uniform refinement was performed for two particles stack. Afterward, particles were combined to do one round ab initio reconstruction. Particles stacks of three classes were refined separately using Non-Uniform refinement and local refinement. Refined particles stack were combined to perform one round ab initio reconstruction. One good class with 959,010 particles was selected. Two rounds of Hetero-refinement were performed. A final 922,712 particles stack was selected to do Non-Uniform refinement and local refinement. Finally, the volume map was refined to 2.86 Å.

For data of S1PR1F, 6,305 micrographs were selected with 5,879,327 particles. Particles extraction and 2D classification were performed the same as S1PR1S. 2D classification and the following 3D refinement were performed in cryoSPARC v3.2.0. 2,670,318 particles were selected after several rounds of 2D classifications. Afterward, ab initio reconstruction was performed. Particles of two good classes with 1,265,555 and 920,714 particles were re-centered and re-extracted separately by Relion with a box size of 288 pixels and a pixel size of 0.83 Å. Meanwhile, another round of ab initio reconstruction was performed separately for the two good classes of the first round ab initio reconstruction. Hetero-refinement was performed using the re-extracted particles and the second round ab initio reconstruction volume as a reference with box size changed to 288 pixels. Particles from three good classes of the two Hetero-refinement results were refined separately using Non-

Uniform refinement and local refinement. The volume map refined from one particles stacks with 909,398 particles reported a higher resolution of 2.83 Å. The particle stacks were selected for further processing. Two rounds of Hetero-refinement were performed. 871,539 particles were selected and further refined using Non-Uniform refinement and local refinement. Moreover, one round of ab initio reconstruction was further performed and 788,201particles of one good class were selected. Non-Uniform refinement and local refinement further refined these particles, resulting in a 2.83 Å map.

For data of S1PR1C, 5,356 micrographs were selected with 4,706,164 particles. Particles extraction was performed the same as the other two datasets. 2D classification and the following 3D refinement were performed in cryoSPARC v.2.14.2. 3,098,124 particles were selected after several rounds of 2D classifications. One round of ab initio reconstruction was performed to further select refined particles stacks with 2,227,503 particles. These particles were re-centered and re-extracted by Relion with a box size of 144 pixels and a pixel size of 1.66 Å. One round of Non-Uniform refinement was performed in cryoSPARC. Particles were re-extracted again by Relion with a box size of 288 pixels and a pixel size of 0.83 Å. Particles were brought back to cryoSPARC. Non-Uniform refinement, local refinement, and ab initio reconstruction were performed. Particles of two good classes were combined and Non-Uniform refinement was performed. Afterward, Hetero-refinement was performed. Particles of two good classes were combined to do Non-Uniform refinement and Local refinement. To obtain a nice particles stack, one round of Hetero-refinement was performed. One class with 592,139 particles was further refined to a 2.98Å map using Non-Uniform refinement and local refinement. Another class with 902,278 particles was also further refined by Non-Uniform refinement and local refinement. Moreover, one round of ab initio reconstruction was performed, yielding 847,759 particles. Non-Uniform and local refinement refined these particles, resulting in a 2.89 Å map. Particle star file conversions between Relion and cryoSPARC were performed mainly using the pyem (7).

#### Model building

The initial template of S1PR1 was generated by the swiss-model (https://swissmodel.expasy.org/), and the initial template of Gαi1, β1, γ2, and scFv16 was obtained from Cryo-EM structure of histamine H1 receptor Gq complex (PDB: 7DFL) (8). These models were docked into the EM density map using dock_in_map in Phenix (9), and followed by iterative rounds of manual building in Coot (10). The final model was subjected to a real space refinement in Phenix (9). The complex model was validated by Molprobity (11, 12). FSC between map and model was calculated by phenix.mtriage (13). Structural figures were prepared in PyMOL (https://pymol.org/2/), UCSF Chimera (14) and UCSF ChimeraX (15).

#### Bioluminescence resonance energy transfer assay (BRET)

S1PR1-mediated $G_{i1}$ protein activation was measured by BRET assays (16). HEK293T cells (ATCC CRL-11268; mycoplasma free) were co-transfected in a 1:1:1:1 ratio of receptor: $G_{\alpha i1}$-Rluc8:$G_{\beta}$:$G_{\gamma}$-GFP2 with polyethyleneimine. After at least 18h, transfected cells were harvested and reseeded in white opaque bottom 96-well assay plates (Beyotime) at a density of 30,000–50,000 cells per well in media (DMEM added 2% dialyzed FBS). The next day, the medium was decanted. Cells were incubated in 40 μL 7.5 μM coelenterazine 400a (Goldbio) in drug buffer (1×Hank's balanced salt solution (HBSS), 20 mM HEPES, pH 7.4) for 2min, and then treated with 20 μL compounds prepared in drug buffer at serial concentration gradient for an additional 5 min. Plates were read in an LB940 Mithras plate reader (Berthold Technologies) with 395-nm and 510-nm emission filters with a 1s per well integration times. BRET ratios were calculated as the ratio of GFP2 emission (510 nm) to Rluc8 emission (395 nm) and analyzed in GraphPad prism 8.0.

#### Molecular dynamics simulations

The Membrane Builder module in CHARMM-GUI server (17) was used to prepare the simulation inputs, including a membrane of pre-equilibrated (310 K) POPC lipids based on the OPM database alignment (18), TIP3P solvent with 0.15 M Na+/Cl- ions, and the CHARMM36 force field (19). The force field of the ligands was generated by the CGenFF program (20).

All MD simulations were performed using GROMACS-2019.4 (21). The CHARMM36 forcefield was used to describe the interactions in the system. Energy minimization was performed for 5000 steps by the steepest descent algorithm. Then a 100 ps NVT simulation was performed at 310 K for solvent equilibration, followed by a 1 ns NPT equilibration to 1 atm using the Berendsen barostat (22). All MD simulations were performed with a time-step of 1 fs. Long-range electrostatic interactions were treated by the particle-mesh Ewald method (23, 24). The short-range electrostatic and van der Waals interactions both used a cutoff of 10 Å. All bonds were constrained by the LINCS algorithm (23, 24). Here, MD simulations were started from the Shallow conformations of the CBP307-bound structure. Simulation is longer than 500 ns and repeated three times.

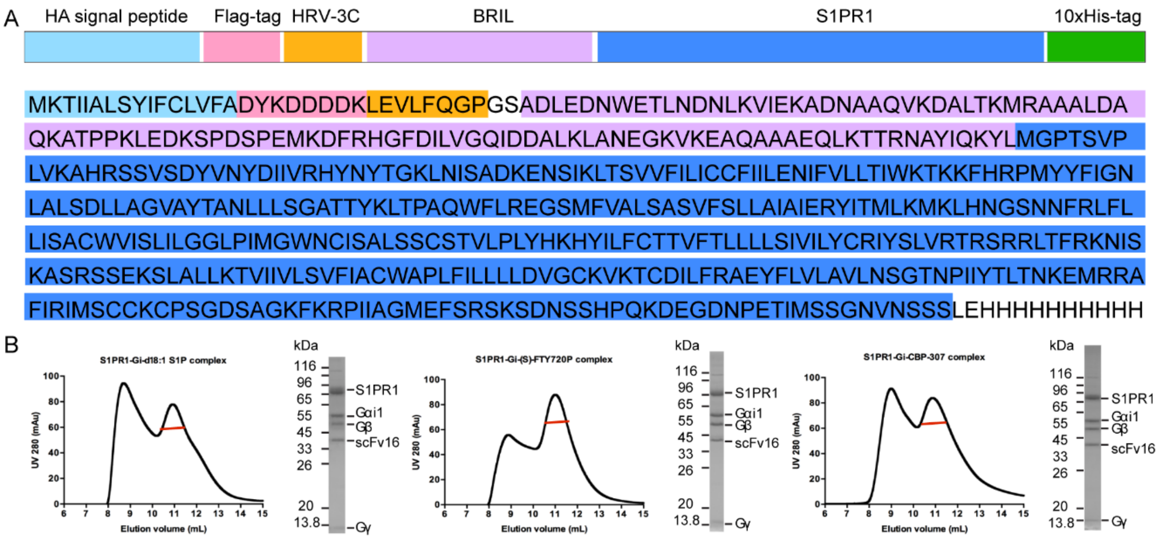

**Figure S1. The amino acid sequence of the S1PR1 construct and chromatography of the S1PR1-G$_i$-scFv16 complex with three small agonists.**

**A**. Sequences are annotated to denote the location of the haemagglutinin (HA) signal sequence (sky blue), Flag tag (pink), HRV-3C protease cleavage sites (bright orange), fusion protein BRIL (violet), S1PR1 (marine), and His tag (green). **B.** Final size exclusion chromatography elution profile of the S1PR1-G$_i$-scFv16 complex with d18:1 S1P, (S)-FTY720-P, and CBP-307. Elution of peak indicated by red line was used for structure determination. SDS–PAGE of one fraction at elution peak is shown, demonstrating the presence of each component of the complex.

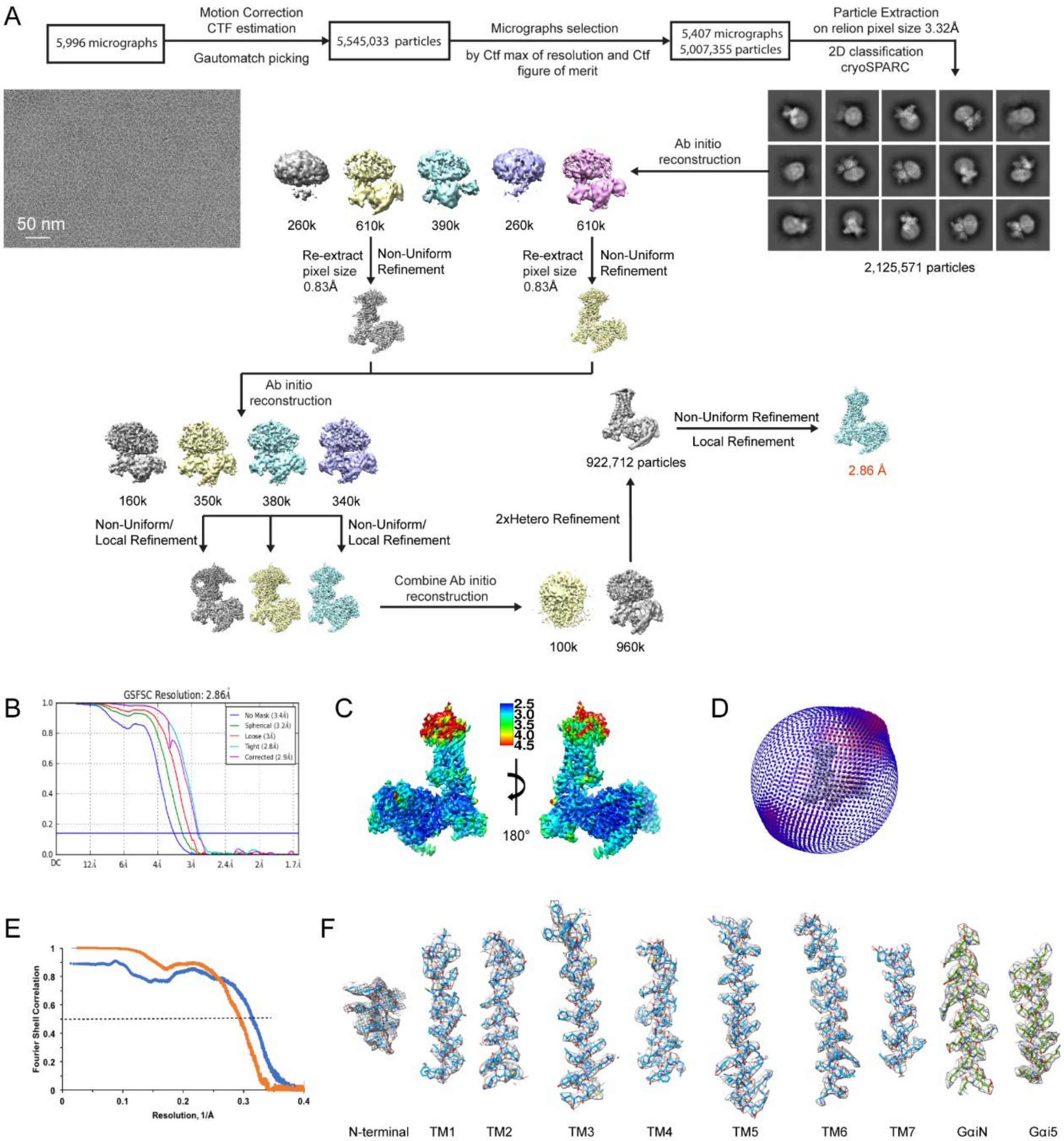

**Figure S2. Cryo-EM reconstruction of S1PR1-$G_i$-scFv16- d18:1 S1P complex.**

**A**. Image processing workflow of S1PR1-$G_i$-scFv16- d18:1 S1P complex. Representative micrograph and 2D classes are shown. **B.** Gold-standard FSC curves used for global-resolution estimates within cryoSPARC. **C.** Local resolution display of the reconstructed 2.86Å map. Local resolution was estimated within cryoSPARC. The resolution range is shown from 2.5 Å-4.5 Å. **D.** Agular distribution of particles used in the final reconstruction of the complex. **E.** FSC curves. FSC curve of the half1 map versus the half2 map (orange). FSC curve of the atomic model against the final map (blue). **F**. Side-chain density of N-terminal and TM helix of S1PR1 as well as $G_{\alpha iN}$ and $G_{\alpha i5}$ helix. The density of S38-L47 on TM1 was missing. Residues shown include N-terminal (D23-N36), TM1 (T48-K72), TM2 (P79-L104), TM3 (P114-L147), TM4 (N157-M180), TM5 (K200-R234), TM6 (E249-G281), TM7 (A293-T314), $G_{\alpha iN}$ (A7-A31) and $G_{\alpha i5}$ (T316-F323). Volume is contoured at a threshold level of 0.15.

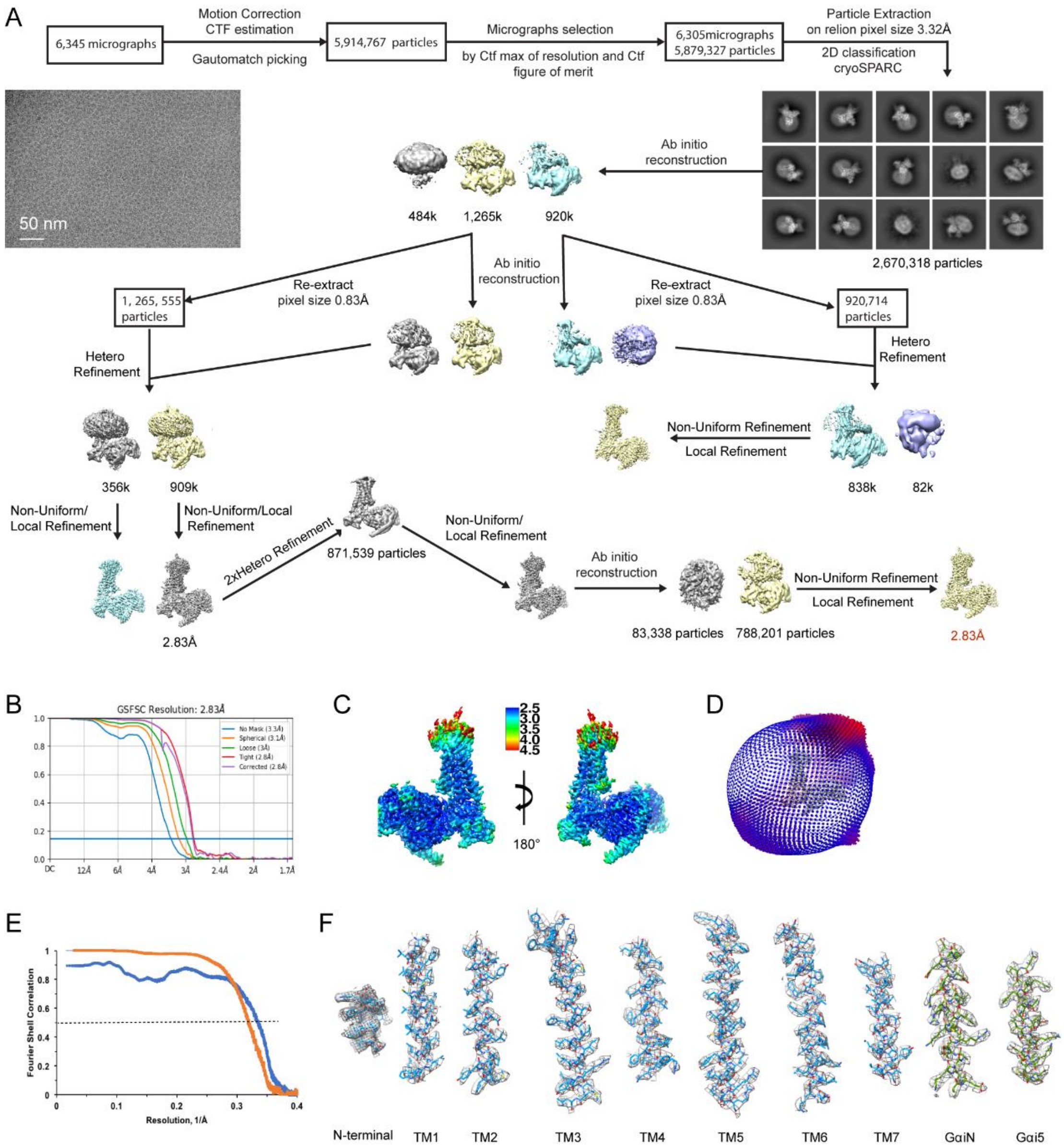

**Figure S3. Cryo-EM reconstruction of S1PR1-G$_i$-scFv16-(S)-FTY720-P complex.**

**A**. Image processing workflow of S1PR1-G$_i$-scFv16-(S)-FTY720-P complex. Representative micrograph and 2D classes were shown. **B.** Gold-standard FSC curves used for global-resolution estimates within cryoSPARC. **C.** Local resolution display of reconstructed 2.83Å map. Local resolution was estimated within cryoSPARC. The resolution range is shown from 2.5 Å-4.5 Å. **D.** Agular distribution of particles used in the final reconstruction of the complex. **E.** FSC curves. FSC curve of the half1 map versus the half2 map (orange). FSC curve of the atomic model against the final map (blue). **F**. Side-chain density of N-terminal and TM helix of S1PR1 as well as G$_{\alpha iN}$ and G$_{\alpha i5}$ helix. The density of S38-L47 on TM1 was missing. Residues shown include: N-terminal (D23-N36), TM1 (T48-K72), TM2 (P79-L104), TM3 (P114-L147), TM4 (N157-M180), TM5 (K200-R234), TM6 (E249-G281), TM7 (A293-T314), G$_{\alpha iN}$ (A7-A31) and G$_{\alpha i5}$ (T316-F323). Volume was contoured at a threshold level of 0.15.

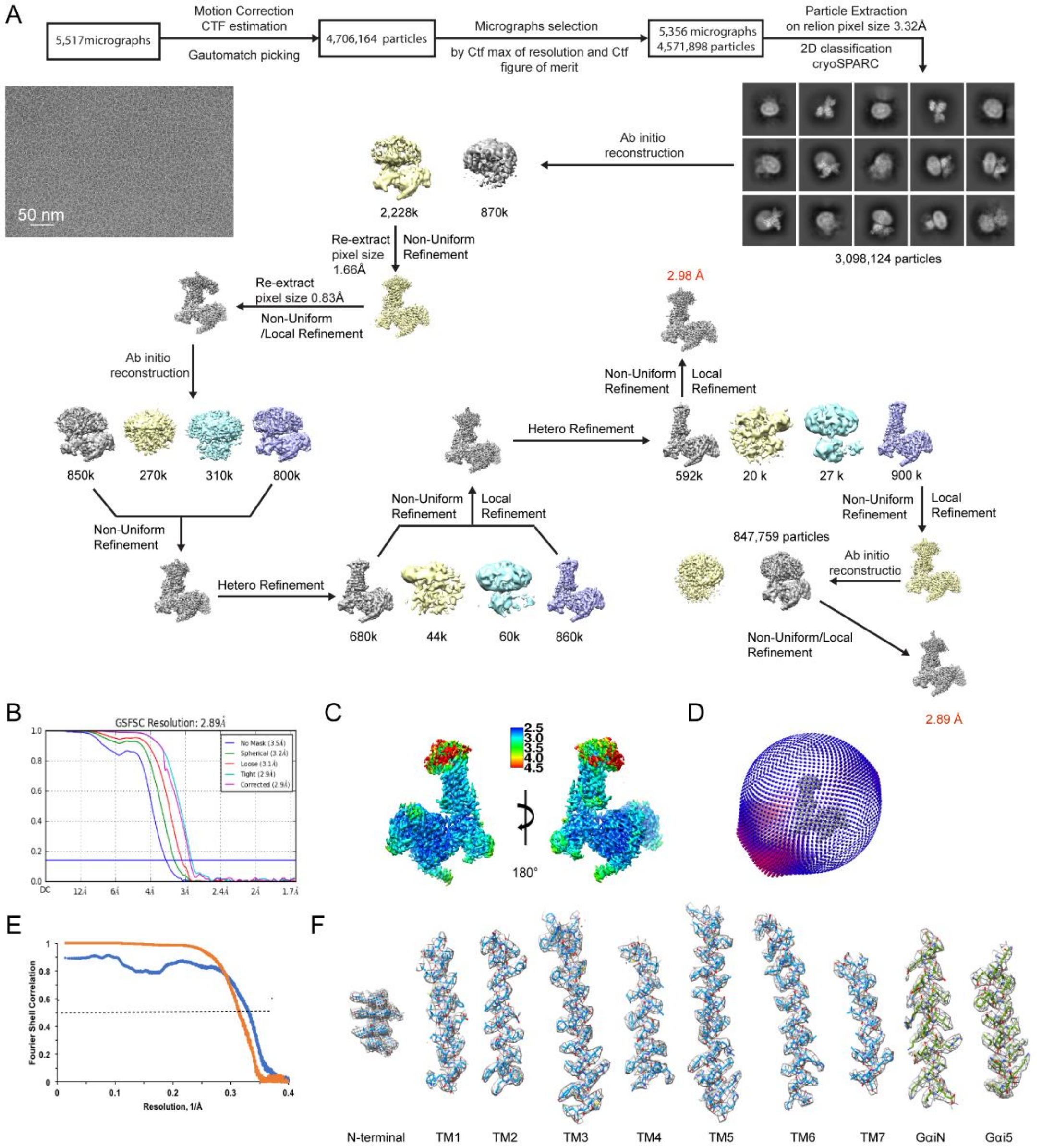

**Figure S4. Cryo-EM reconstruction of S1PR1-$G_i$-scFv16-CBP-307 complex.**

**A**. Image processing workflow S1PR1-$G_i$-scFv16-CBP-307 complex. Representative micrograph and 2D classes were shown. **B.** Gold-standard FSC curves of 2.89 Å map used for global-resolution estimates within cryoSPARC. **C.** Local resolution display of the reconstructed 2.89 Å map. Local resolution of 2.89 Å map was estimated within cryoSPARC. The resolution range is shown from 2.5 Å-4.5 Å. **D.** Agular distribution of particles used in the final reconstruction of the 2.89 Å map. **E.** FSC curves between model and map as well as between half map for the model of 2.89 Å map. FSC curve of the half1 map versus the half2 map (orange). FSC curve of the atomic model against the final map (blue). **F**. Side-chain density of N-terminal and TM helix of S1PR1 as well as $G_{\alpha iN}$ and $G_{\alpha i5}$ helix of the 2.89 Å map. The density of S38-L47 on TM1 was missing. Residue shown include: N-terminal (D23-N36), TM1 (T48-K72), TM2 (P79-L104), TM3 (P114-L147), TM4 (N157-M180), TM5 (K200-R234), TM6 (E249-G281), TM7 (A293-T314), $G_{\alpha iN}$ (A7-A31) and $G_{\alpha i5}$ (T316-F323). Volume was contoured at a threshold level of 0.15.

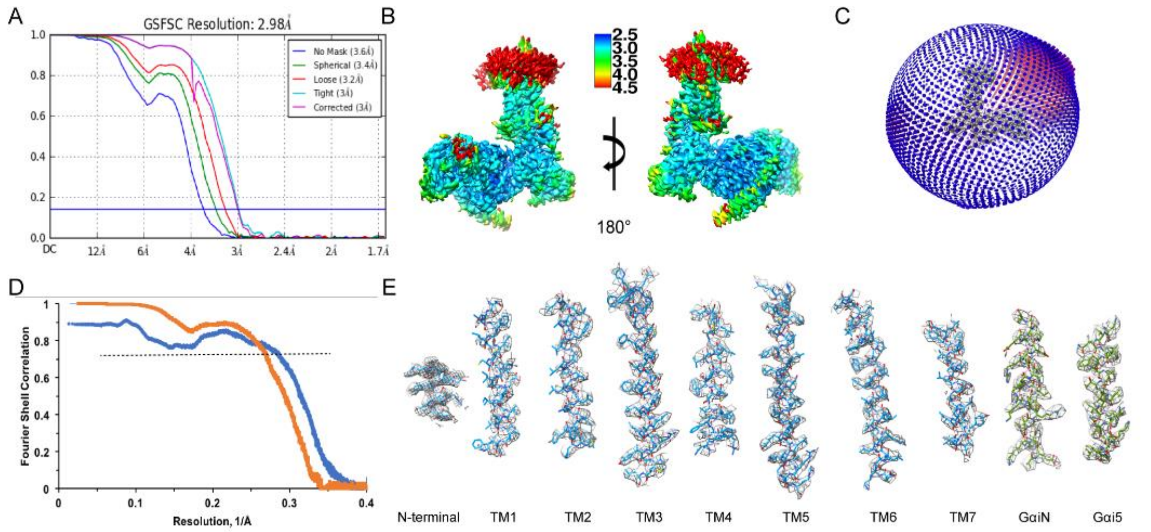

**Figure S5. FSC curves, local resolution, angular distribution, and side-chain density of 2.98 Å map of S1PR1-G$_i$-scFv16-CBP-307 complex.**

**A.** Gold-standard FSC curves of 2.98 Å map used for global-resolution estimates within cryoSPARC. **B.** Local resolution display of the reconstructed 2.98 Å map. Local resolution of 2.98 Å map was estimated within cryoSPARC. The resolution range is shown from 2.5 Å-4.5 Å. **C.** Agular distribution of particles used in the final reconstruction of the 2.98 Å map. **D.** FSC curves between model and map as well as between half map for the model of the 2.98 Å map. FSC curve of the half1 map versus the half2 map (orange). FSC curve of the atomic model against the final map (blue). **E**. Side-chain density of N-terminal and TM helix of S1PR1 as well as $G_{\alpha iN}$ and $G_{\alpha i5}$ helix of the 2.98 Å map. The density of S38-L47 on TM1 was missing. Residue shown include: N-terminal (D23-L35), TM1 (T48-K72), TM2 (P79-L104), TM3 (P114-L147), TM4 (N157-M180), TM5 (K200-R234), TM6 (E249-G281), TM7 (A293-T314), $G_{\alpha iN}$ (A7-A31) and $G_{\alpha i5}$ (T316-F323). Volume was contoured at a threshold level of 0.15.

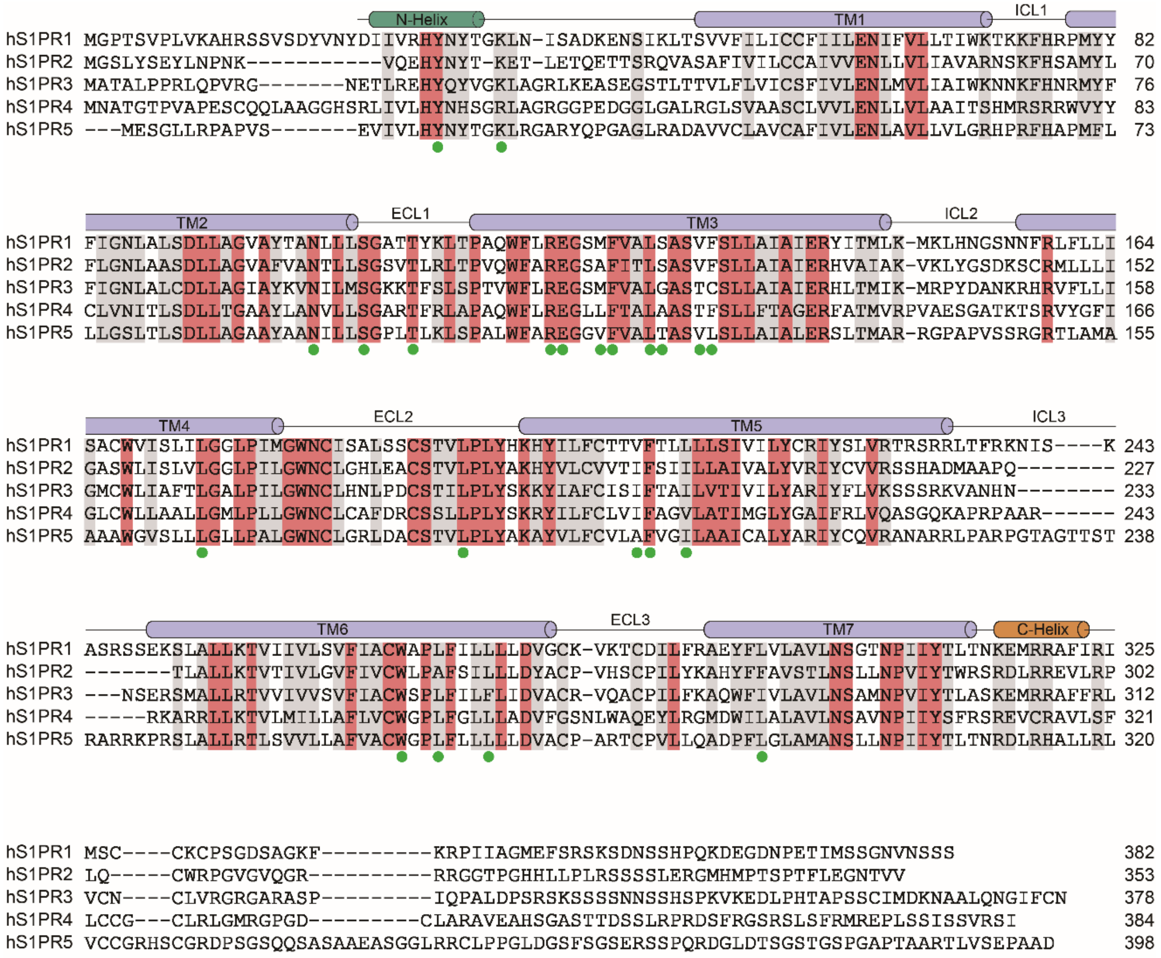

**Figure S6. Sequence alignment of human S1PR1, S1PR2, S1PR3, S1PR4, and S1PR5**

Secondary structural elements of human S1PR1 are displayed above the alignment. Invariant and highly conserved residues are shaded deep salmon and gray, respectively. The d18:1 S1P binding pocket residues are indicated by the green circle below.

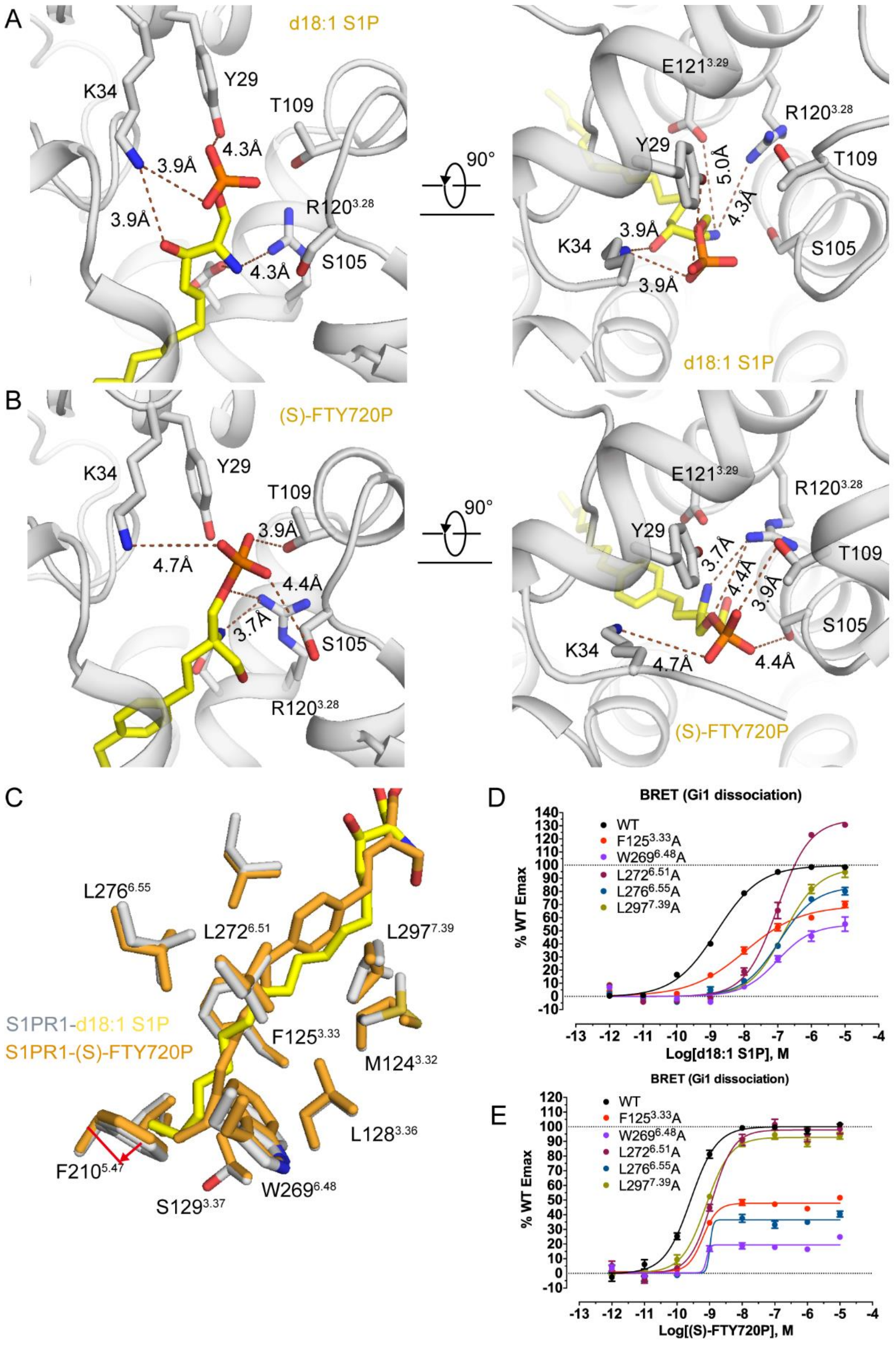

**Figure S7. Comparison of d18:1 S1P and (S)-FTY720-P binding S1PR1.**

**A.** Interaction between the polar groups of d18:1 S1P and S1PR1. d18:1 S1P and key residues are shown in sticks. Dashed lines indicate the interactions with distance. Side and top views are shown. **B.** Interaction between the polar groups of (S)-FTY720-P and S1PR1. (S)-FTY720-P and key residues are shown in sticks. Dashed lines indicate the interactions with distance. Side and top views are shown. **C.** Superposition of hydrophobic residues around the acyl chain of d18:1 S1P (yellow) and (S)-FTY720-P (orange), including L195$^{ECL2}$, L276$^{5.55}$, F210$^{5.47}$, S129$^{3.37}$, W269$^{5.48}$, L128$^{3.36}$, M124$^{3.32}$, F125$^{3.33}$, L272$^{5.51,}$ and L297$^{7.39}$, shown in sticks. **D.** The effects of mutants W269$^{5.48}$A, L272$^{5.51}$A and L276$^{5.55}$A of S1PR1 on d18:1 S1P induced $G_i$ signal activation measured by $G_i$ dissociation assay (BERT assay). All data are mean ± SEM of three independent experiments for wild-type or mutants. **E.** The effects of mutants W269$^{5.48}$A, L272$^{5.51}$A, and L276$^{5.55}$A of S1PR1 on (S)-FTY720-P induced $G_i$ signal activation measured by $G_i$ dissociation assay (BERT assay). All data are mean ± SEM of three independent experiments for wild-type or mutants.

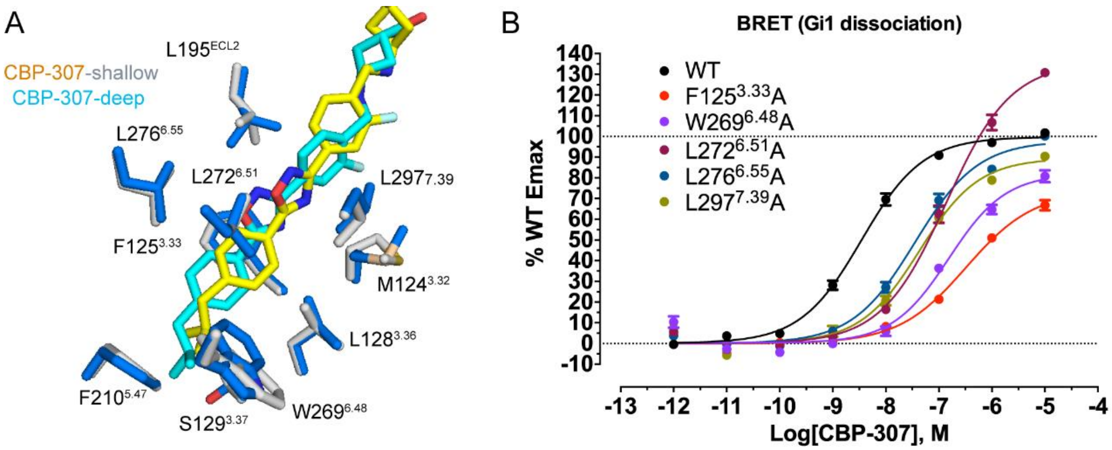

**Figure S8. CBP-307 interaction with hydrophobic residues in S1PR1 ligand-binding pocket.**

**A.** Superposition of hydrophobic residues around the hydrophobic tail of CBP-307 in two binding modes. CBP-307 (yellow) with S1PR1 (gray) in shallow binding mode and CBP-307 (marine) with S1PR1 (marine) in deep binding mode. Residues including L195$^{ECL2}$, L276$^{5.55}$, F210$^{5.47}$, S129$^{3.37}$, W269$^{5.48}$, L128$^{3.36}$, M124$^{3.32}$, F125$^{3.33}$, L272$^{5.51}$ and L297$^{7.39}$, are shown in sticks. **B.** The effects of W269$^{5.48}$A, L272$^{5.51}$A, and L276$^{5.55}$A mutations of S1PR1 on CBP-307 induced $G_i$ signal activation measured by $G_i$ dissociation assay (BERT assay). All data are mean ± SEM of three independent experiments for wild-type or mutants.

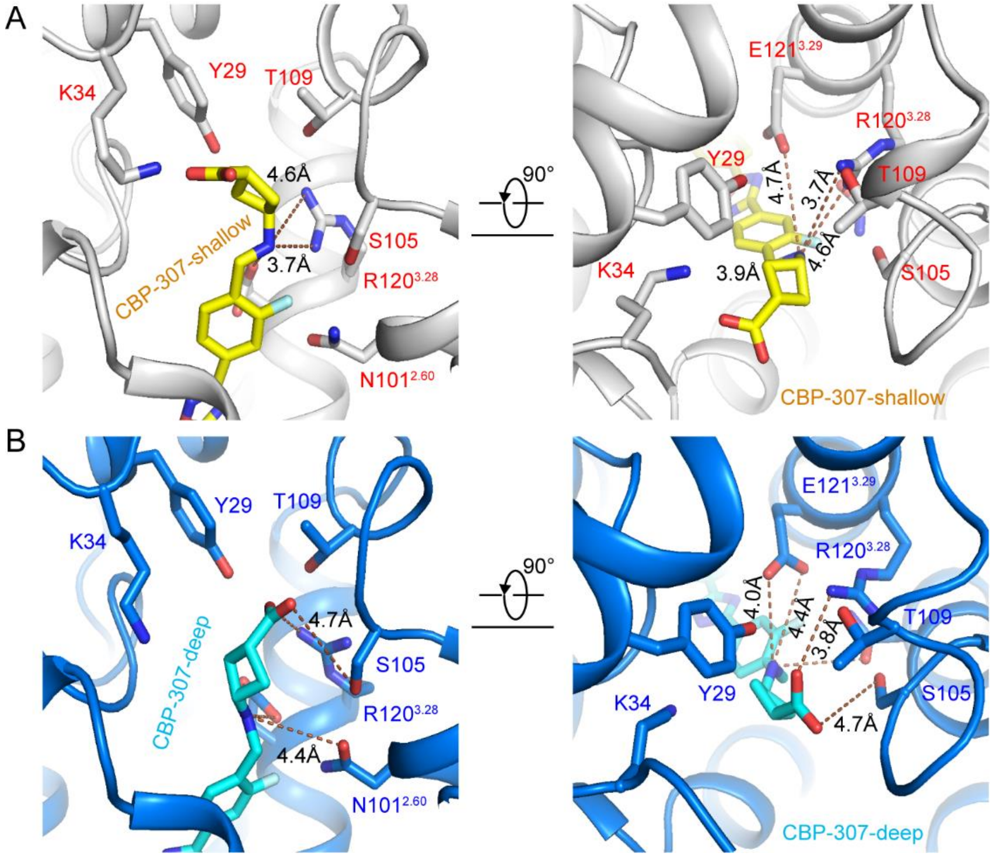

**Figure S9. Comparison of other weak and indirect interactions in the two binding modes of CBP-307 bound S1PR1.**

**A.** Weak or indirect interactions between the polar groups of CBP-307 and S1PR1 in the shallow binding mode. CBP-307 and key residues are shown in sticks. Dashed lines indicate weak or indirect interactions with distance. Side and top views are shown. **B.** Weak and indirect interactions between the polar groups of CBP-307 and S1PR1 in the deep binding mode. CBP-307 and key residues are shown in sticks. Dashed lines indicate weak or indirect interactions with distance. Side and top views are shown.

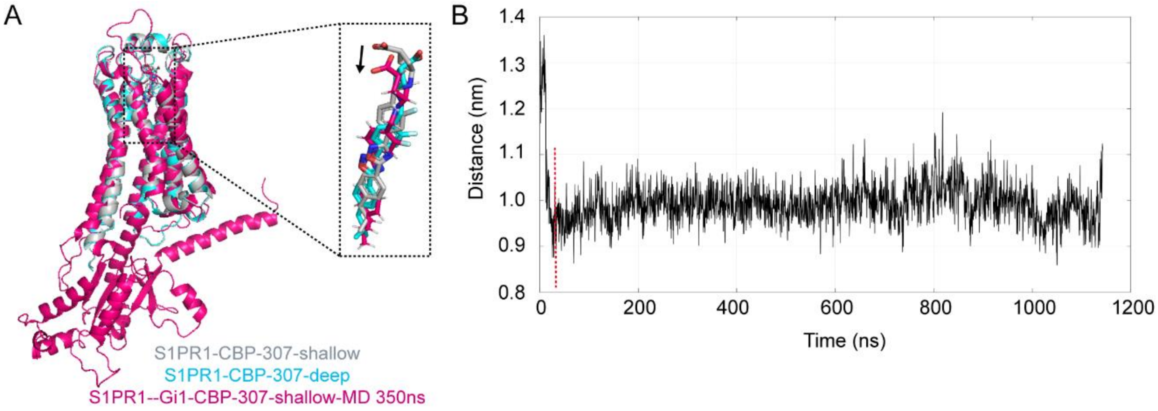

**Figure S10. Molecular dynamics simulations of CBP-307 bound S1PR1 with $G_i$ in shallow mode.**

A. Comparison of MD at 350ns of CBP-307 bound S1PR1 with $G_i$ in shallow mode (hot pink) with CBP-307 bound S1PR1 in shallow mode (gray) and deep mode (cyan). CBP-307 is shown in sticks. The magnified view of the alignment of CBP-307 in three structures is shown in the inset. B. The distance between atom carbon-05 of CBP-307, the bottom carbon atom at the bottom ring of CBP-307, and the $C_\alpha$ atom of L213 in the MD simulation of CBP-307 bound S1PR1 with $G_i$ in shallow mode. L213 is located at the bottom of the ligand-binding pocket of CBP-307 bound S1PR1.

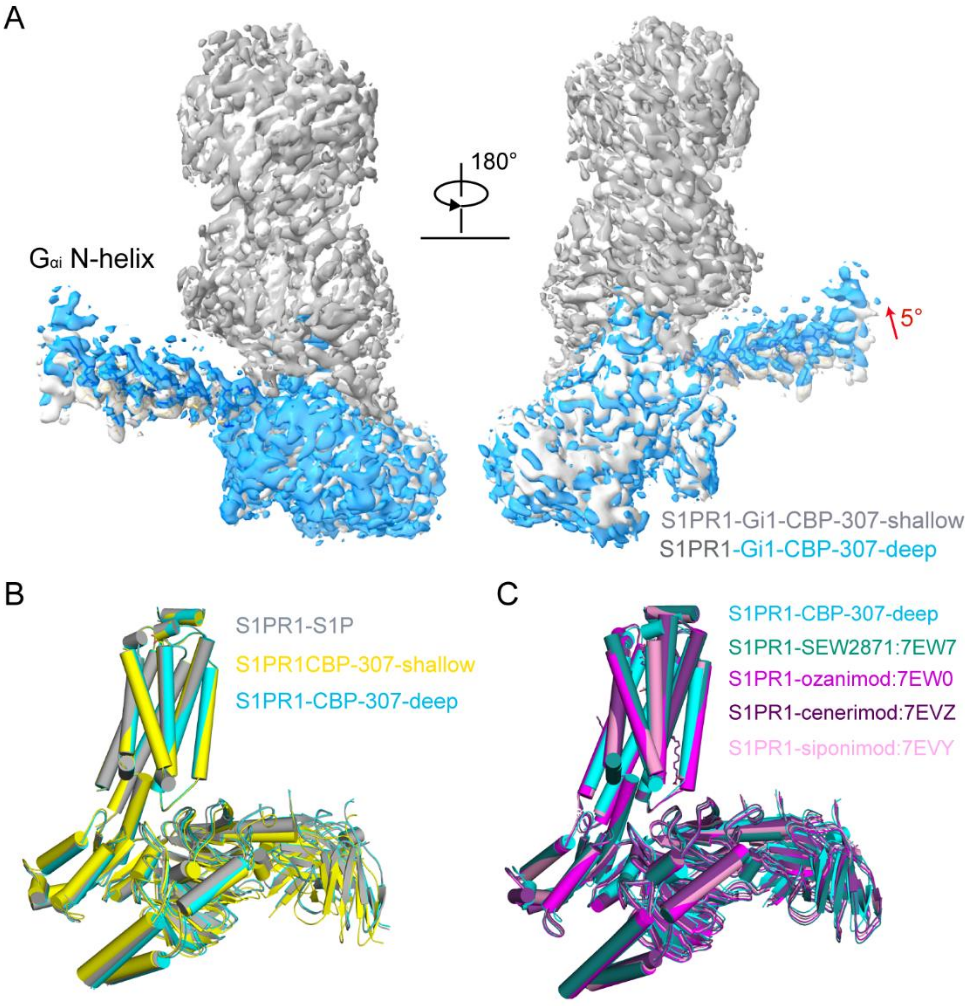

**Figure S11. The density maps of CBP-307 bound S1PR1 and the structural comparison of reported representative S1PR1 structures.**

**A.** The densities of the N-terminal helix of $G_{\alpha i}$ in two CBP-307 binding modes are aligned together. Alignment is performed only at the S1PR1 receptor. Densities of S1PR1 in two CBP-307 binding modes are colored in light gray (Shallow mode) and gray (deep mode), respectively. The densities of the N-terminal helix of $G_{\alpha i}$ in two CBP-307 binding modes are colored in light gray (Shallow mode) and marine (deep mode), respectively. Residues at the N-terminal helix of $G_{\alpha i}$ in two CBP-307 binding modes are shown as sticks and colored in light gray (Shallow mode) and blue (deep mode), respectively. The densities of the N-terminal helix of $G_{\alpha i}$ in two CBP-307 binding modes are rotated. **B.** Superposition of d18:1 S1P bound S1PR1-$G_i$ complex structure (gray) with CBP-307 bound two binding modes of S1PR1-$G_i$ complex structures (shallow: yellow; deep: cyan)**. C.** Superposition of CBP-307 bound S1PR1-$G_i$ complex structures in deep mode (cyan) with four other agonists bound S1PR1-$G_i$ complex structures determined by Shao group: SEW2971 bound (PDB: 7EW7, green), ozanimod bound (PDB: 7EW0, violet), cenerimod bound (PDB: 7EVZ, violet purple), Siponimod bound (PDB: 7EVY, pink)**.**

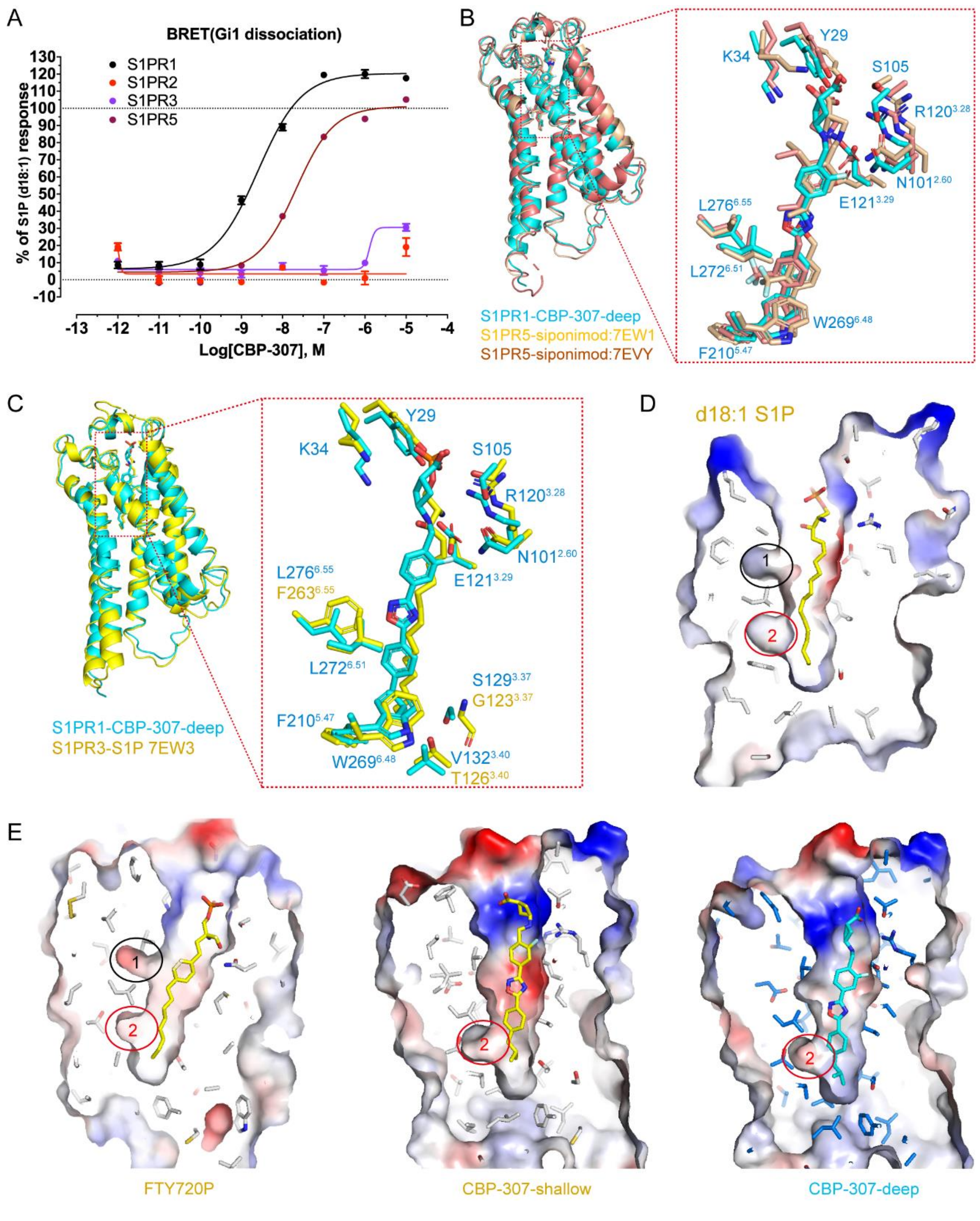

**Figure S12. Receptor selectivity of CBP-307, comparison of representative S1PR1/3/5 structures, and sub-pockets of S1PR1 in d18:1 S1P, (S)-FTY720-P and CBP-307 bound structures.**

**A.** CBP-307 induced $G_i$ signal for S1PRs measured by $G_i$ dissociation assay. All data are mean ± SEM of three independent experiments. **B.** Comparison of CBP-307 bound S1PR1 structure in deep mode (cyan) with Siponimod bound S1PR1 (PDB: 7EVY, rose) and S1PR5 structures (PDB: 7EW1, wheat). A magnified view of agonists and key residues surrounded is shown in the inset. Agonists and residues of S1PR are shown in sticks. **C.** Comparison of CBP-

307 bound S1PR1 structure in deep mode (cyan) with d18:1 S1P bound S1PR3 structure (PDB: 7EW3, yellow). A magnified view of agonists and key residues surrounded is shown in the inset. Agonists and residues of S1PR1 and S1PR3 are shown in sticks. **D.** A cutaway representation of d18:1 S1P binding pocket and two unoccupied sub-pockets in d18:1 S1P bound S1PR1 structure. **E.** Cutaway represents agonist binding pockets and unoccupied sub-pockets in (S)-FTY720-P and CBP-307 bound S1PR1 structures.

**Table S1. Cryo-EM data collection, refinement, and validation statistics**

| | S1PR1-d18:1 S1P (EMDB-32006) (PDB 7VIE) | S1PR1-(S)-FTY720-P (EMDB-32007) (PDB 7VIF) | S1PR1-CBP-307 (Deep) (EMDB-32008) (PDB 7VIG) | S1PR1-CBP-307 (Shallow) (EMDB-32009) (PDB 7VIH) |
|---|---|---|---|---|
| **Data collection and processing** | | | | |
| Magnification | 105,000 | 105,000 | 105,000 | 105,000 |
| Voltage (kV) | 300 | 300 | 300 | 300 |
| Electron exposure (e–/Å$^2$) | 56.14-66.50 | 56.2 | 56.61 | 56.61 |
| Defocus range (μm) | -1.1 - -1.9 | -1.1 - -1.9 | -1.1 - -1.9 | -1.1 - -1.9 |
| Pixel size (Å) | 0.83 | 0.83 | 0.83 | 0.83 |
| Symmetry imposed | C1 | C1 | C1 | C1 |
| Initial particle images (no.) | 5,007,355 | 5,879,327 | 4,571,898 | 4,571,898 |
| Final particle images (no.) | 922,712 | 788,201 | 847,759 | 592139 |
| Map resolution (Å) | 2.86 | 2.83 | 2.89 | 2.98 |
| FSC threshold | 0.143 | 0.143 | 0.143 | 0.143 |
| Map resolution range (Å) | 2.5-4.5 | 2.5-4.5 | 2.5-4.5 | 2.5-4.5 |
| **Refinement** | | | | |
| Initial model used (PDB code) | 7DFL/7C4S | 7DF1/7C4S | 7DF1/7C4S | 7DF1/7C4S |
| Model resolution (Å) | 3.03 | 2.98 | 3.02 | 3.18 |
| FSC threshold | 0.5 | 0.5 | 0.5 | 0.5 |
| Model resolution range (Å) | | | | |
| Map sharpening *B* factor (Å$^2$) | | | | |
| Model composition | | | | |
| Non-hydrogen atoms | 8905 | 8918 | 8917 | 8924 |
| Protein residues | 1134 | 1135 | 1133 | 1136 |
| Ligands | 1 | 1 | 1 | 1 |
| *B* factors (Å$^2$) | | | | |
| Protein | 46.48 | 49.61 | 38.17 | 69.68 |
| Ligand | 73.68 | 67.02 | 50.31 | 98.39 |
| R.m.s. deviations | | | | |
| Bond lengths (Å) | 0.003 | 0.004 | 0.004 | 0.004 |
| Bond angles (°) | 0.605 | 0.609 | 0.651 | 0.775 |
| Validation | | | | |
| MolProbity score | 1.46 | 1.45 | 1.56 | 1.7 |
| Clashscore | 8.59 | 7.98 | 7.64 | 10.17 |
| Ramachandran plot | | | | |
| Favored (%) | 98.11 | 97.94 | 97.21 | 97.05 |
| Allowed (%) | 1.89 | 2.06 | 2.79 | 2.95 |
| Disallowed (%) | 0 | 0 | 0 | 0 |

**Table S2. Statistics of $G_i$ dissociation of S1PR1 mutants**

| S1PR1 variants | d18:1 S1P | | CBP-307 | | (S)-FTY720-P | |
|---|---|---|---|---|---|---|
| | $EC_{50}$, nM ($pEC_{50}$ ± SEM) | Fold | $EC_{50}$, nM ($pEC_{50}$ ± SEM) | Fold | $EC_{50}$, nM ($pEC_{50}$ ± SEM) | Fold |
| WT | 1.62 (8.79 ± 0.11) | 1.00 | 3.55 (8.45 ± 0.09) | 1.00 | 0.23 (9.64 ± 0.08) | 1.00 |
| Y29A | 56.23 (7.25 ± 0.03) | 34.71 | 48.98 (7.31 ± 0.19) | 13.80 | 0.51 (9.29 ± 0.11) | 2.22 |
| K34A | 56.23 (7.25 ± 0.06) | 34.71 | 6.03 (8.22 ± 0.17) | 1.70 | 0.56 (9.25 ± 0.07) | 2.43 |
| N101A | 467.74 (6.33 ± 0.11) | 288.73 | 10.72 (7.97 ± 0.13) | 3.02 | 61.66 (7.21 ± 0.18) | 268.09 |
| S105A | 199.53 (6.70 ± 0.04) | 123.17 | 1.07 (8.97 ± 0.09) | 0.30 | 3.80 (8.42 ± 0.05) | 16.52 |
| T109A | 87.10 (7.06 ± 0.07) | 53.77 | 2.57 (8.59 ± 0.11) | 0.72 | 0.79 (9.10 ± 0.03) | 3.43 |
| R120A | 53.70 (7.27 ± 0.23) | 33.15 | 17.38 (7.76 ± 0.13) | 4.90 | 1737.80 (5.76 ± 0.32) | 7555.65 |
| E121A | 371.54 (6.43 ± 0.19) | 229.35 | 3090.30 (5.51 ± 0.45) | 870.51 | 691.83 (6.16 ± 0.39) | 3007.96 |
| M124A | 45.71 (7.34 ± 0.11) | 28.22 | 114.82 (6.94 ± 0.20) | 32.34 | 1.70 (8.77 ± 0.03) | 7.39 |
| F125A | 11.22 (7.95 ± 0.08) | 6.93 | 323.59 (6.49 ± 0.01) | 91.15 | 0.60 (9.22 ± 0.08) | 2.61 |
| L128A | 11.22 (7.95 ± 0.05) | 6.93 | 34.67 (7.46 ± 0.10) | 9.77 | 0.81 (9.09 ± 0.07) | 3.52 |
| S129A | 7.94 (8.10 ± 0.10) | 4.90 | 2.51 (8.60 ± 0.11) | 0.71 | 0.17 (9.77 ± 0.16) | 0.74 |
| V132A | 1.51 (8.82 ± 0.13) | 0.93 | 8.13 (8.09 ± 0.29) | 2.29 | 0.16 (9.79 ± 0.12) | 0.70 |
| F133A | 6.76 (8.17 ± 0.03) | 4.17 | 15.85 (7.80 ± 0.37) | 4.46 | 0.19 (9.72 ± 0.07) | 0.83 |
| L174A | 7.94 (8.10 ± 0.07) | 4.90 | 11.75 (7.93 ± 0.39) | 3.31 | 0.29 (9.54 ± 0.09) | 1.26 |
| L195A | 28.18 (7.55 ± 0.03) | 17.40 | 48.98 (7.31 ± 0.15) | 13.80 | 0.47 (9.33 ± 0.03) | 2.04 |
| V209A | 6.76 (8.17 ± 0.08) | 4.17 | 8.71 (8.06 ± 0.27) | 2.45 | 0.30 (9.52 ± 0.07) | 1.30 |
| F210A | 21.88 | 13.51 | 3.98 | 1.12 | 0.12 | 0.52 |

| | | | | | | |
|---|---|---|---|---|---|---|
| | (7.66 ± 0.18) | | (8.40 ± 0.19) | | (9.91 ± 0.04) | |
| L213A | 0.54<br>(9.27 ± 0.37) | 0.33 | 3.16<br>(8.59 ± 0.18) | 0.89 | 0.34<br>(9.47 ± 0.17) | 1.48 |
| W269A | 81.28<br>(7.09 ± 0.21) | 50.17 | 147.91<br>(6.83 ± 0.08) | 41.66 | 0.4<br>(9.40 ± 0.24) | 1.74 |
| L272A | 93.33<br>(7.03 ± 0.11) | 57.61 | 138.04<br>(6.86 ± 0.12) | 38.88 | 1.15<br>(8.94 ± 0.02) | 5.00 |
| L276A | 109.65<br>(6.96 ± 0.06) | 67.69 | 35.48<br>(7.45 ± 0.08) | 9.99 | 1.20<br>(8.92 ± 0.06) | 5.22 |
| L297A | 151.36<br>(6.82 ± 0.06) | 93.43 | 42.66<br>(7.37 ± 0.07) | 12.02 | 0.74<br>(9.13 ± 0.07) | 3.22 |
| G106A | 33.11<br>(7.48 ± 0.07) | 20.44 | 2.51<br>(8.60 ± 0.11) | 0.71 | 0.25<br>(9.60 ± 0.12) | 1.09 |
| Y110A | 10.23<br>(7.99 ± 0.08) | 6.31 | 7.24<br>(8.14 ± 0.09) | 2.04 | 0.33<br>(9.48 ± 0.08) | 1.43 |
| W117A | 147.91<br>(6.83 ± 0.09) | 91.30 | 25.70<br>(7.59 ± 0.08) | 7.24 | 0.93<br>(9.03 ± 0.04) | 4.04 |
| N36A | 4.68<br>(8.33 ± 0.10) | 2.89 | 2.34<br>(8.63 ± 0.09) | 0.66 | 0.28<br>(9.55 ± 0.05) | 1.22 |
| E294A | 25.70<br>(7.59 ± 0.10) | 15.86 | 6.46<br>(8.19 ± 0.08) | 1.82 | 0.20<br>(9.69 ± 0.07) | 0.87 |

All data are the mean ± SEM (n = 3 independent experiments). Fold: the EC50 of mutant relative to the EC50 of WT.
